# Dynamical forces drive cell and organ morphology changes during embryonic development

**DOI:** 10.1101/2024.07.13.603371

**Authors:** Raj Kumar Manna, Emma M. Retzlaff, Anna Maria Hinman, Yiling Lan, Osama Abdel-Razek, Mike Bates, Heidi Hehnly, Jeffrey D. Amack, M. Lisa Manning

## Abstract

Cells, tissues, and organs must change shape in precise ways during embryonic development to execute their functions. Multiple mechanisms including biochemical signaling pathways and biophysical forces help drive these morphology changes, but it has been difficult to tease apart their contributions, especially from tissue-scale dynamic forces that are typically ignored. We use a combination of mathematical models and *in vivo* experiments to study a simple organ in the zebrafish embryo called Kupffer’s vesicle. Modeling indicates that dynamic forces generated by tissue movements in the embryo produce shape changes in Kupffer’s vesicle that are observed during development. Laser ablations in the zebrafish embryo that alter these forces result in altered organ shapes matching model predictions. These results demonstrate that dynamic forces sculpt cell and organ shape during embryo development.

**Significance Statement:** We aim to understand the mechanisms that control precise cell and tissue shape changes required for organ function. Many studies focused on cell shapes have ignored the role of dynamic forces self-generated by slow tissue flows, but recent work showing tissues are near a jamming transition with diverging relaxation timescales suggests slow motion could give rise to large forces. Through a combination of mathematical modeling, imaging, and mechanical perturbations to *in vivo* experiments, our work demonstrates that tissue-scale dynamic forces are sculpting the shape of an epithelial organ in the zebrafish embryo called Kupffer’s vesicle (KV). Because there are many processes during development that occur at similarly slow rates, this suggests that self-generated dynamic forces should be investigated more broadly.

## Main text

A key question in developmental biology is how organisms robustly control the morphology of cells, tissues, and organs,^1^ given that information for such structures is ultimately encoded at the scale of molecules. Mechanical forces, such as drag forces generated by viscous dissipation, or tension on individual cell-cell interfaces could help provide such control mechanisms. Because tissue motion is so slow, many past studies focused on cell shapes and other small-scale tissue features have assumed local force balance and ignored dynamical forces like drag.^2-9^ In more recent work, researchers have relaxed this condition, allowing frictional or drag forces at the boundary of a tissue to explain tissue morphodynamics.^10, 11^ But viscous forces are not necessarily limited to the boundaries; large-scale hydrodynamic models for tissues capture bulk tissue flows that contribute to large-scale tissue morphology changes.^12, 13^ Therefore, an open question is whether cell-cell interactions can self-generate large-scale tissue flows, resulting in dynamical forces that feed back onto smaller-scale features, ultimately controlling cell and organ shapes.

In this work, we demonstrate that self-generated bulk dynamical forces -- those that arise fromcells and tissues moving at a finite rate during embryo development -- control cell and organ shape in Kupffer’s Vesicle (KV), the organ that establishes the left-right (LR) body axis in zebrafish. This highlights that slow rates of tissue-scale motion can generate large cell-scale forces if the tissue relaxation timescales are also large, and suggests that self-generated dynamic forces are likely governing cell and tissue morphology in other developmental processes.

We focus here on KV as a simple, small model organ that is composed of a single layer of monociliated epithelial cells surrounding a fluid-filled lumen that develops in the tailbud of the zebrafish embryo (Fig 1A-C).^14-16^ Other vertebrates have a similar ciliated organ that controls LR patterning.^17-20^ In zebrafish, motile cilia generate a counterclockwise fluid flow and associated shear stresses inside the lumen, which are sensed on the left-anterior side of the organ and trigger downstream signals that orient the LR axis relative to the anterior-posterior (AP) axis of the embryo.^21-24^

**Fig. 1.**
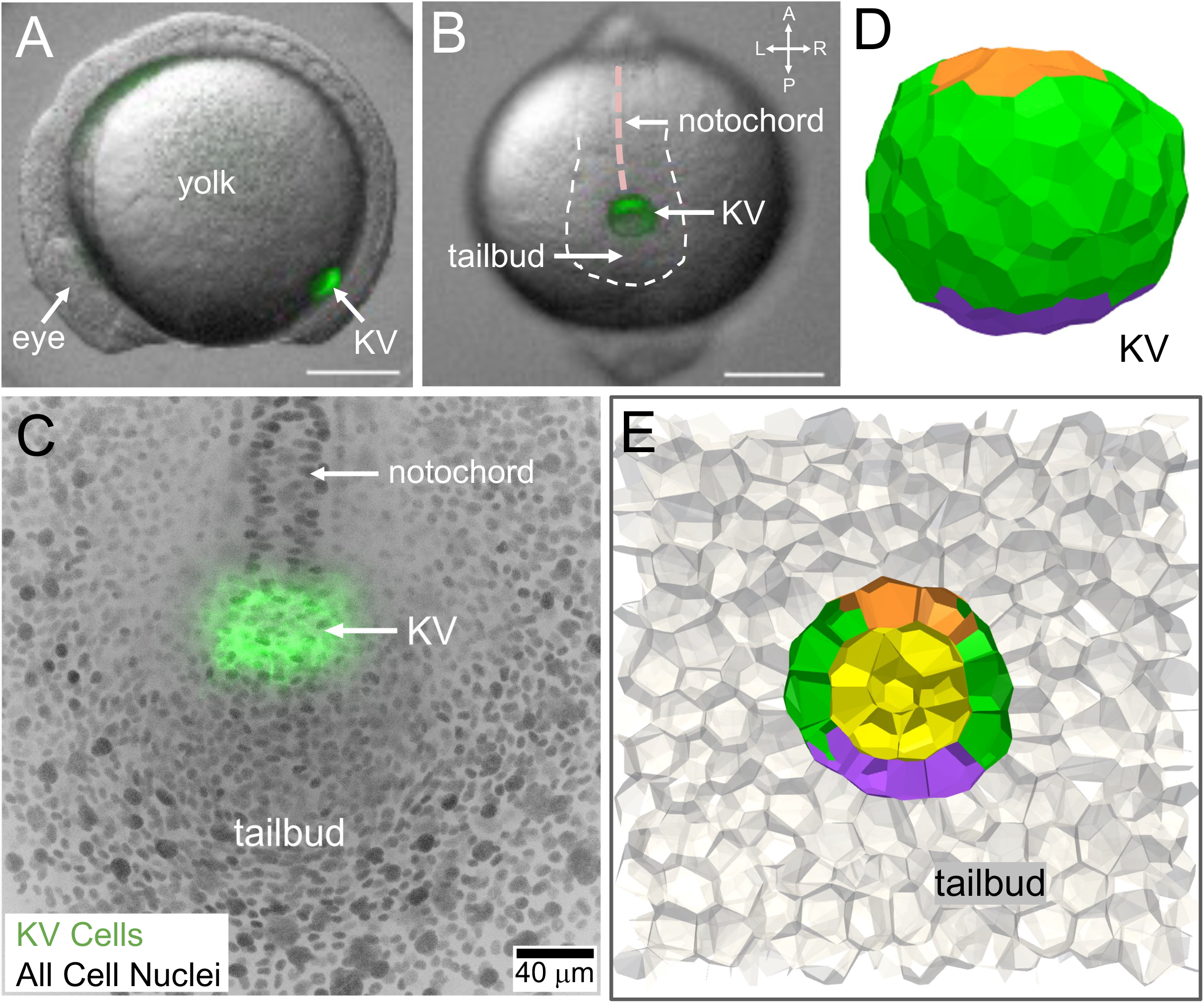
Zebrafish Kupffer’s Vesicle (KV) as a model organ for investigating shape changes during organogenesis. (**A-B**) KV marked by GFP expression (green) in a living zebrafish embryo is located at the posterior end of the notochord in the tailbud seen in the lateral view (A) and dorsal view (B) at the 12-somite stage. Scalebar = 200 μm. A = anterior, P = posterior, L = left, R = right. Dotted pink line demarcates the notochord. Dotted white lines outline the approximate boundary of the tailbud. (**C**) Zoom in view of a 3D projection of the KV and tailbud tissue at the KV dorsal side obtained from a confocal microscope (scalebar = 20 μm). (**D**) KV is modeled as one-layer of cells on the surface of a sphere. Anterior and posterior part of the KV is marked respectively by orange and purple colors. (**E**) The middle plane of KV along with tailbud cells (marked by grey color).

In previous work, some of us have identified a series of cell shape changes during KV morphogenesis, termed ‘KV remodeling,’ that are required for KV function.^25^ In KV remodeling, anterior cells elongate to allow a denser packing of cells and cilia, while cells on the posterior stretch in the opposite direction to reduce cell/cilia density. We and others have identified perturbations that prevent this remodeling; these perturbations also reduce the rate and directionality of fluid flow across the anterior of the lumen and interfere with correct LR patterning.^24, 26, 27^

What mechanisms generate cell shape changes during KV remodeling? No morphogen gradients have been implicated despite many searches, and lumen expansion is also not required.^23^ Recent work has focused on physical interactions between the notochord, which is located just to the anterior of KV inside the embryo and undergoes convergent extension during KV remodeling. Researchers have found evidence that direct physical interactions between KV cells and the extracellular matrix that is adjacent to the notochord are contributing to cell shape change.^27^ This would correspond to a *static* gradient in the mechanical environment around KV. There is also a static gradient in tissue stiffness in the zebrafish tailbud around the KV, which may also impact cell shape,^28^ though initial modeling suggests it may not be sufficient.^25^

We hypothesize that since the KV is dynamically moving through surrounding tissue precisely during the window of KV remodeling, perhaps that motion could generate a *dynamic* gradient in the mechanical environment that leads to changes in cell and organ shape. Initially, this seems implausible, as the typical rate of KV motion is about a micron a minute, which is quite slow and similar to rates of cell motion in other development processes. Because of these slow rates, researchers typically assume the system is force balanced at each timestep (e.g. no dynamic gradient) in both experiments (e.g. in stress monolayer microscopy^7^ or video force microscopy^6, 9^) and in mathematical models (e.g. vertex models^2, 4^).

However, recent work has also demonstrated that many tissues, including the zebrafish tailbud, operate close to a fluid-solid transition.^28^ Near such transitions, the effective viscosity, or equivalently the material relaxation timescale, diverges.^29^ In these cases, very slow motions can give rise to large forces. Previous work on oocyte development in Drosophila^30^ has suggested that viscous drag forces help determine the shape of an egg. Moreover, at low Reynolds number (i.e. high viscosity), the drag forces that KV would experience should be similar to a sphere moving through a viscous fluid, which would compress the posterior side of the organ (and cells that comprise it) and extend the anterior side of the organ. Initial simulations of a simple, spherically symmetric mathematical model for the KV geometry in 2D^31^ and 3D^32^ indicated that dynamical drag forces could plausibly generate the observed shape changes.

To directly test our hypothesis, we need a model that accounts for realistic, asymmetric forces that drive the KV through the tissue in control experiments, and then we need to perturb the rate at which KV moves through the surrounding tissue in both model and experiment. By developing novel 3D models and laser ablation experiments, we 1) quantify how perturbing structures in and around the KV impacts KV motion, 2) measure the shape of the organ in these perturbed cases, and 3) demonstrate that observed shape changes *in vivo* are precisely those predicted to result from the altered dynamic forces.

### KV moves through the tailbud tissue with a well-defined speed and lumen shape

To visualize and quantify development of the KV organ in the context of its surrounding cellular environment, we used double stable transgenic *Tg(sox17:EGFP-CAAX,myl7:EGFP);Tg(ubb:LOXP-mScarlet-NLS-LOXP-QF-GAL4)* embryos–referred to here as *Tg(Sox17:EGFP-CAAX);Tg(ubi:mScarlet-NLS)* embryos–that express membrane-localized EGFP (green fluorescent protein) in KV cells and nuclear-localized mScarlet (red fluorescent protein) in all cells including tailbud cells that surround KV.^33^ Capturing images of these live embryos every two minutes on a spinning disk confocal microscope allows for the analysis of cell behaviors and cell-cell interactions in the tailbud during KV morphogenesis (Movie S1).

Live imaging of wild-type embryos was used to analyze the movement of the KV through the surrounding tailbud tissue during KV remodeling that occurs between the three-somite stage (3ss) and seven-somite stage (7ss) (2-hour window) of development (Movie S2). We use particle image velocimetry to quantify the bulk motions of the tissue. Using previously validated methods ^32^, we quantify the tissue velocity fields at the KV midplane relative to the frame where the KV is at rest (Methods; Fig S17A), and then compute the average velocity of the tailbud tissue in the AP direction at different distances from the KV (Fig. S17B). Consistent with previous work ^32^, we find a strong gradient in tissue velocities in the surrounding tailbud, indicating that bulk, tissue-scale dynamic drag forces are acting on the KV. At the resolution necessary to see global tissue movements, it is difficult to precisely resolve the shapes of individual cells, so we largely focus on the shape of the KV lumen as a robust readout of these dynamic forces. For each timepoint, we trace the lumen shape at the maximum cross-section of the KV (Methods; Fig. S1) and compute the centroid and dimensionless shape parameter (*Sr*) – the ratio of y- and x-components of the radius of gyration (Methods; Fig. S6), with *Sr* =1 for a sphere, *Sr* < 1 for an oblate spheroid, and *Sr* > 1 for a prolate spheroid. We compute the average KV speed *u* from the motion of the lumen centroid between frames. In control wildtype embryos (n=9) KV moves at a well-defined speed *u* that is on the order of 1 micron/minute and increases with increasing somite stage, with a twofold increase between 3ss and 7ss (Fig. 2G, Fig. S5A). The KV also has a well-defined oblate shape with *Sr <* 1 in control embryos at these developmental stages (Fig. 2H, Fig. S10A).

**Fig. 2.**
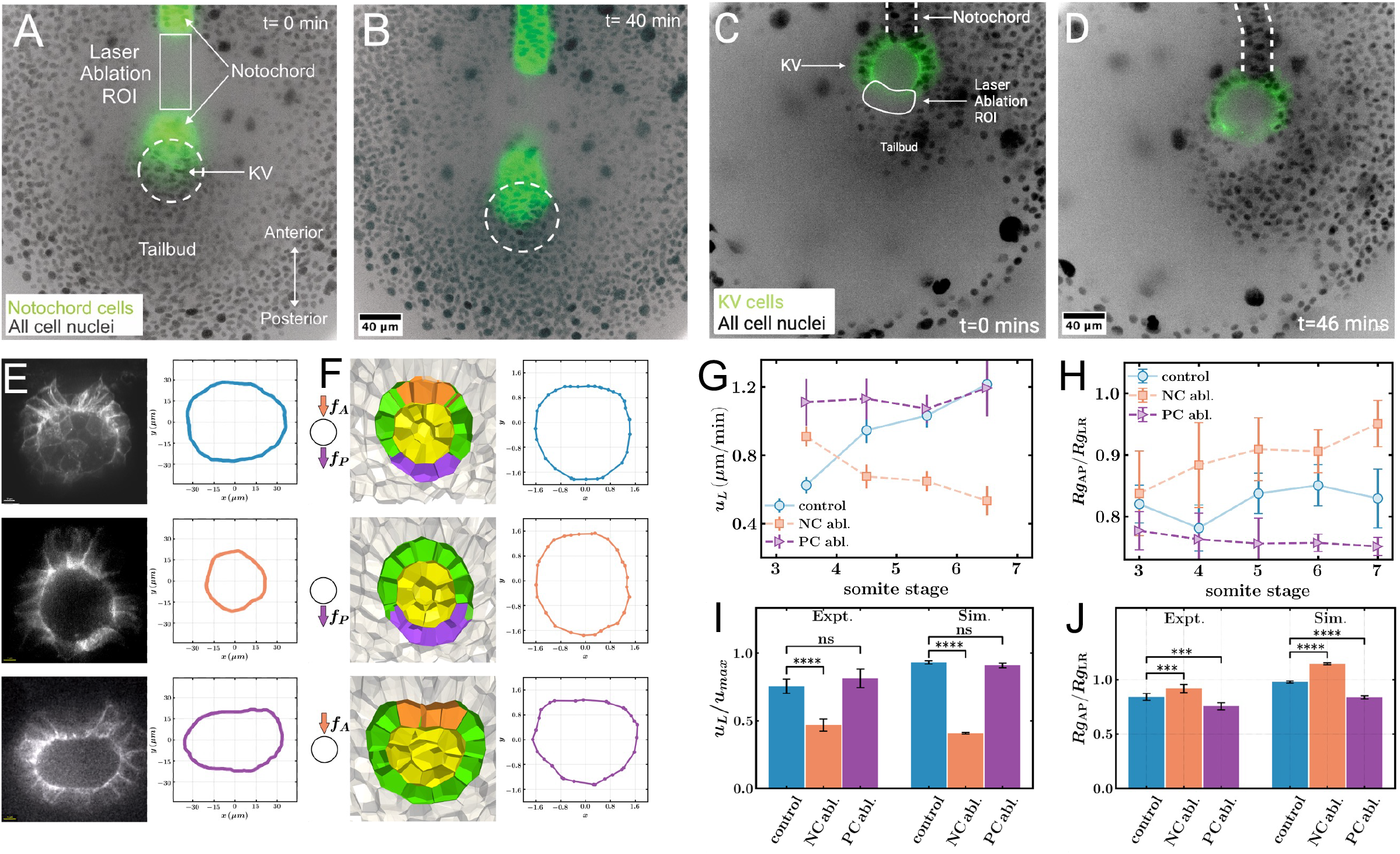
Ablation perturbations indicate that changes to dynamic forces alter the KV cell shapes. (**A-D**) Microscopy images of laser ablations, with all cell nuclei labeled in black. (A-B) Notochord ablation at the time of ablation, t=0, (A) and 40 minutes post-ablation (B). Notochord cells are labeled green, and the location of KV is highlighted with dashed white circle. The white box indicates the ablated region of interest (ROI). (C-D) Posterior cell ablation at the time of ablation, t=0, (C) and 46 minutes post-ablation (D). KV cell membranes are labeled green, and the location of the notochord is indicated with dashed white lines. (**E**) Confocal microscopy images (left) and traces of lumen shapes (right) for control (top) notochord ablated (middle) and posterior cell ablated embryos (bottom). (**F**) Snapshots from simulations (left) and traces of lumen shapes (right) for simulations with both pushing from the notochord and pulling from posterior cells (top), simulation of notochord ablation with only posterior cells pulling (middle), simulation of posterior ablation with only notochord cells pushing (bottom). (**G-J**) Quantification of speed of lumen and shape of lumen for different conditions. (G) Experimental data for the average speed of the center of the lumen (u_L_) as a function of somite stage. (H) Experimentally observed lumen shape quantified as the ratio of the radius of gyration of lumen along the AP axis (Rg_AP_) to that along the LR axis (Rg_LR_). (I) The experimental lumen speed measurements averaged over 5-7ss for different experimental conditions compared to their simulation counterparts. (J) The experimental lumen shape measurements averaged over 5-7ss compared to simulation counterparts. Error bars represent standard error.

### Models with anterior and posterior forces recapitulate wildtype KV motion and lumen shape

Previous work has demonstrated strong, posterior-directed cell migration for cells in the posterior of the tailbud.^34^ We hypothesize this process generates pulling forces originating in the posterior region of KV, and that there are also pushing forces from the anterior driven by convergent extension in the notochord. To model the impact of these forces on KV, we simulate a 3D vertex model^35^ with multiple tissue types,^36^ including KV cells embedded within a three-dimensional tailbud tissue (Fig. 1D,E, see Methods), and we again quantify KV speed and lumen shape changes, and KV cell shape changes (Fig S11). We identify a parameter regime where the 3D model with both types of forces can quantitatively recapitulate both the speed of the lumen and the oblate lumen shape of KV (Top panel of Fig. 2F, and Fig. 2I,J), as well as cell shape changes (Fig 3E). In the simulations, the oblate shape results from an interplay of anterior pushing forces, posterior pulling forces, and the resistance of the viscoelastic tailbud tissue surrounding KV, as shown in Supplemental Movie S8. In these simulations where the KV reaches a steady state velocity, the viscous stresses are precisely balancing the active pushing and pulling forces. Since this argument is very general, we expect that the result is independent of the details of the model. To demonstrate this, we confirm that these simulation results are robust to changes to vertex model parameters such as the magnitude of active forces (Fig. S12) and heterotypic interfacial tension (Fig. S18), as well as KV and tailbud fluidity (Fig. S18). In addition, we also simulate a much simpler hydrodynamic model of a membrane surrounded by a highly viscous medium, with anterior pushing forces and posterior pulling forces (see Methods, Movie S11). We find that the shape of the membrane is quite similar to both the KV lumen and the lumen in the full 3D vertex model (Fig S12).

**Fig. 3.**
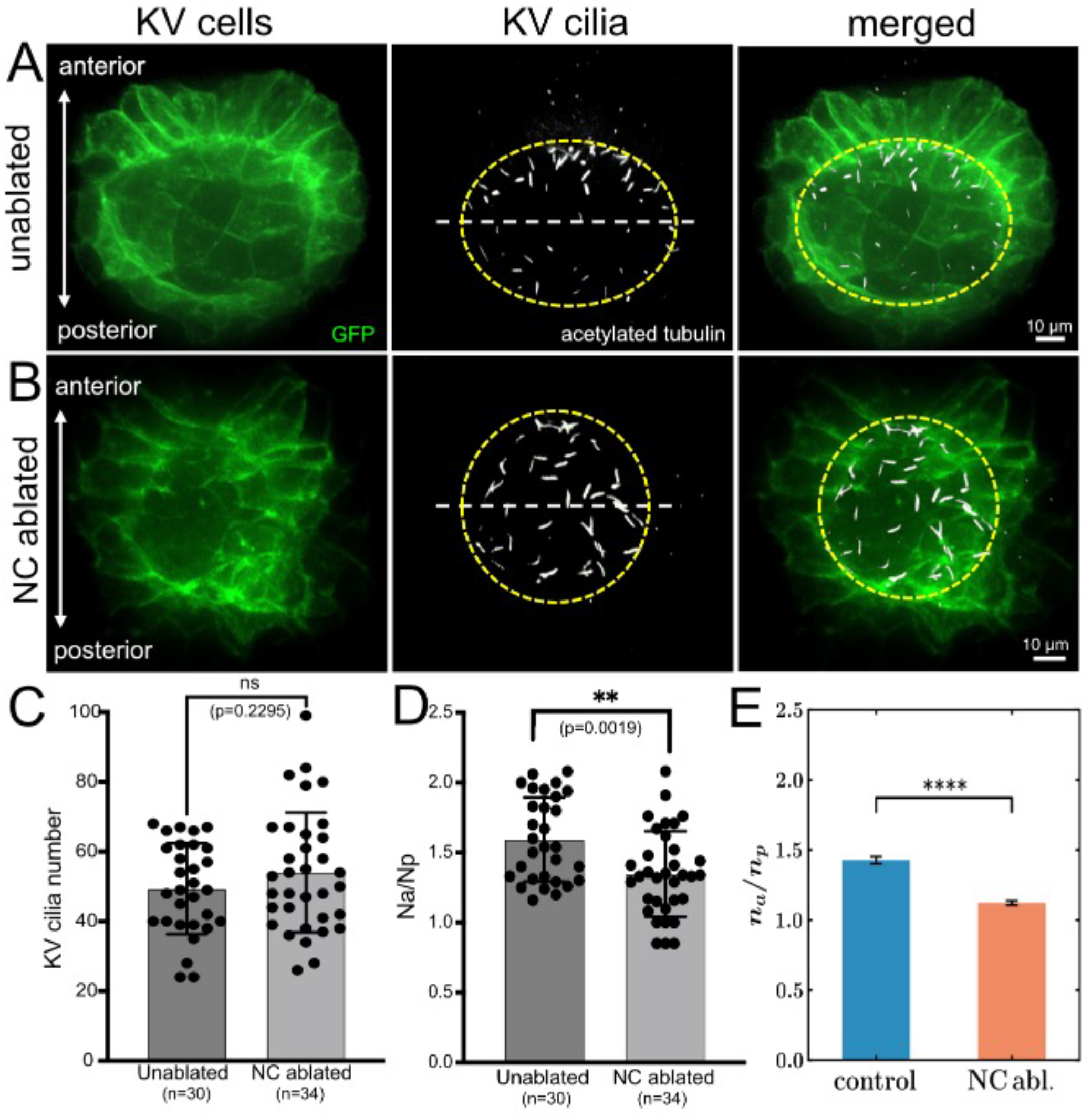
AP distribution ciliated KV cells is disrupted by notochord ablation. (**A-B**) Microscopy images of KV in an unablated control embryo (A) and notochord (NC) ablated embryo (B)with GFP-labeled KV cell membranes in green (left), cilia labeled with acetylated tubulin antibodies in white (middle), and merged images (right). a. Yellow dashed line approximates the KV lumen boundary and the white dashed line bisects KV into anterior and posterior regions. (**C**) The total number of KV cilia observed in unablated and NC ablated embryos. Each point represents an embryo. (**D**) The ratio of the number of anterior cilia (Na) to the number of posterior cilia (Np) for unablated and NC ablated embryos indicates reduced AP asymmetry of cilia and less KV cell shape remodeling in NC ablated embryos. n=number of embryos analyzed. Error bars=1 standard deviation. ns=not significant, **p<0.01 P values are indicated in each graph. (**E**) The ratio of the number of anterior KV cells to the number of posterior KV cells in control and NC ablation cases obtained from simulation.

### Ablation experiments to perturb dynamic forces

We next devised laser ablation experiments to directly perturb the forces exerted on the KV within live embryos without directly perturbing biochemical signaling pathways. To disrupt pushing forces generated by the rod-like notochord on the KV, we used a pulsed laser to ablate a defined section of the notochord just anterior to the KV (Fig. 2A-B, Movie S3). We use *Tg(−2*.*4shha-ABC:GFP)* transgenic embryos that express GFP in notochord cells^37^ to define the region to ablate and assess ablation efficacy. The laser-induced lesion in the notochord persisted throughout live imaging of KV remodeling (2 hours) and was still apparent 24 hours post-ablation (Fig S13). These results indicate laser ablation provides an effective approach to mechanically perturb the notochord. In addition to a notochord pushing force acting on KV, live imaging reveals that posterior KV cells are highly protrusive (Movie S4), which is reminiscent of leader cells during collective migration.^38^ To disrupt pulling forces these posterior cells may have on KV, we ablated a subset of posterior cells in the middle plane of KV in *Tg(Sox17:EGFP-CAAX);Tg(ubi:mScarlet-NLS)* embryos (Fig. 2C-D, Movie S5). Ablating 3 to 8 posterior cells did not disrupt the KV lumen. Consistent with ablating only a small number of cells, no discernable defects were observed 24 hours post-ablation (Fig S14).

### Ablation of notochord reduces the speed of KV

Laser ablation of the notochord at ∼3ss significantly reduces the movement of the KV, as illustrated in Fig. 2B (snapshot taken 40 minutes post-ablation) and Movie S6. Using the same technique to measure KV speed as for wildtype embryos, we found that, in contrast to wildtype, the average speed of KV in the notochord ablation experiments (n=7) decreases between 3ss and 7ss (Fig. 2G, Fig. S2, Fig. S5B). For notochord ablations, the average speed of KV at the 7-somite stage is half of that observed in the control experiments (Fig. 2G,), and significantly reduced as compared to controls when averaged over 5ss to 7ss (Fig. 2I). Control ablation of cells in the head region of the embryo far away from notochord and KV (n=6) have no effect on KV speed (Fig. S4, Fig. S5D, Fig. S15A,B) or morphogenesis (Fig. S9, Fig. S10D, Fig. S15C,D). To mimic the effect of notochord ablation in our 3D vertex model, we investigate KV dynamics when the anterior pushing forces vanish (*f*_*a*_ *= 0*), while the posterior forces are still acting (middle panel of Fig. 2F). In our overdamped limit, the velocity of KV is directly proportional to the force acting on it, and so we can fit the observed experimental KV speed data by assuming that the anterior notochord pushing provides half the total force on the KV. This demonstrates that forces in the notochord region generate KV movement towards the posterior.

### Notochord ablation alters global morphology of KV by elongating along the AP axis

Next, we use our 3D vertex model to predict how the KV organ will change shape in the presence of these perturbed dynamic forces (Movie S9). In contrast to wildtype lumens that are oblate, our model predicts that after the notochord pushing forces are set to zero, the KV and lumen shape should continuously extend along the AP axis so that the shape becomes more prolate, and Sr increases (Fig. 2F,J). This is precisely what we observe in the notochord ablation experiments -- after ablation, the lumen shape undergoes elongation along the anterior-posterior axis (Fig. 2E). While in control experiments, the shape parameter remains relatively constant at a value less than 1, notochord ablation significantly increases the average lumen shape (Fig. 2H,J, Fig. S7, Fig. S10B). Thus, forces originating from notochord extension are not only crucial for KV motion but also serve as vital factors in maintaining the physiological lumen shape observed in control experiments.

### Laser ablation of a small region of KV posterior cells does not affect the speed of KV

Because notochord ablation does not completely halt KV movement, there must be other forces acting on KV. We hypothesized that active pulling forces from posterior cells represent one such mechanism. To study this, we selectively laser-ablate a small cluster of posterior cells at the middle plane of KV. We were limited to the smaller ablation region to ensure we could preserve the integrity of the fluid-filled lumen (see Fig. 2C and Methods for details). We found that this small perturbation to posterior cells did not impede KV speed (Movie S7); tracked KV speed post-ablation (n=7) was consistent with that observed in control embryos (Fig. 2G,I, Fig. S3, Fig. S5C). We speculate that perhaps we were only able to disrupt a subset of the cells pulling downward, due to our necessarily limited ablation region, and therefore this perturbation was not sufficient to reduce KV speed.

### Posterior cells ablation elongates the KV along the LR axis

Although our posterior ablation was not sufficient to change KV speed, given that notochord pushing and tissue drag forces are still acting, we hypothesize that the change in location of the forces could still be sufficient to alter the KV shape. To simulate this ablation, we set pulling forces to zero (*f*_*p*_ *= 0*) and increase pushing force to match the observed KV speed in the experiment (Movie S10). Our analysis reveals that the shape parameter for the lumen is lower than that of control experiments, indicating elongation along the left-right axis and a shift towards an oblate shape compared to the control condition (Fig. 2F,J). Examination of lumen shape in embryos subjected to posterior cell ablation (n=7) is more oblate than controls (Fig. 2E, Fig. S8, Fig. S10C) and shows that the average shape parameter remains consistently lower than in control experiments and is statistically significant between the 5-7 ss (Fig. 2H,J), as predicted by our 3D vertex model. Therefore, our finding suggests the location of dynamic forces is pivotal for the physiological shape changes of KV observed during development.

### KV cell shapes and Anterior/Posterior cilia distribution are altered in notochord ablated embryos, but asymmetry and LR patterning persist

Since changes to dynamical forces are sufficient to alter lumen shape in ablated embryos, we wonder if these dynamical forces also change the individual cell shapes to drive KV remodeling. We first focus on the anterior/posterior (AP) distribution of cilia, which is related to cell shape since each cell is monocilated. Wild-type embryos exhibit a highly asymmetric AP distribution of cilia, with ∼60% of ciliated cells positioned in the anterior half of KV.^26, 37, 39-41^ We use the number of ciliated cells in the anterior region divided by the number in the posterior region, Na/Np, to describe this AP asymmetry. As shown in Fig 3A-D, notochord ablation does not alter the total number of ciliated KV cells but significantly reduces the AP distribution (Na/Np) as compared to unablated controls (see also Table S1). The 3D vertex model also predicts this reduction of AP asymmetry due to notochord ablation (Fig. 3E). To directly analyze KV cells shapes, we calculated a length-to-width ratio (LWR) of anterior and posterior KV cells in which the apical width and cell length can be clearly resolved as we have previously described^25, 26, 41^. The average LWR of anterior cells and posterior cells was not significantly different between NC ablated embryos and unablated controls, which is likely due to high variability in overall cell shapes. We next focused on apical width of KV cells (Fig 4A,B). Mechanistically, a reduced Na/Np could result from a defect in anterior KV cells failing to reduce their apical surfaces to allow tight packing of ciliated cells and/or a defect in posterior KV cells increasing apical surfaces to reduce cilia density in the posterior region. Since this analysis did not depend on resolving KV cell length, we could include more cells. The average apical width of anterior KV cells was significantly increased in notochord ablated embryos as compared to unabated controls (Student’s t-test p=0.0141) (Fig. 4C). There was also a trend of a reduced apical width in posterior KV cells in notochord ablated embryos, but this was not statistically significant (Student’s t-test p=0.1725) (Fig. 4D). Similar changes to the cell shapes are also observed in 3D vertex models (Fig. 4E,F). This is consistent with our interpretation that dynamic forces contribute to the remodeling of KV cell shapes, in addition to the overall organ shape. However, this perturbation is not enough to destroy the anterior/posterior asymmetry -- there is still a higher fraction of ciliated cells in the anterior region than in the posterior in ablation experiments (Table S1). Moreover, LR patterning and heart laterality is unaffected in these embryos (Fig S16; Table S2; Table S3).

**Fig. 4.**
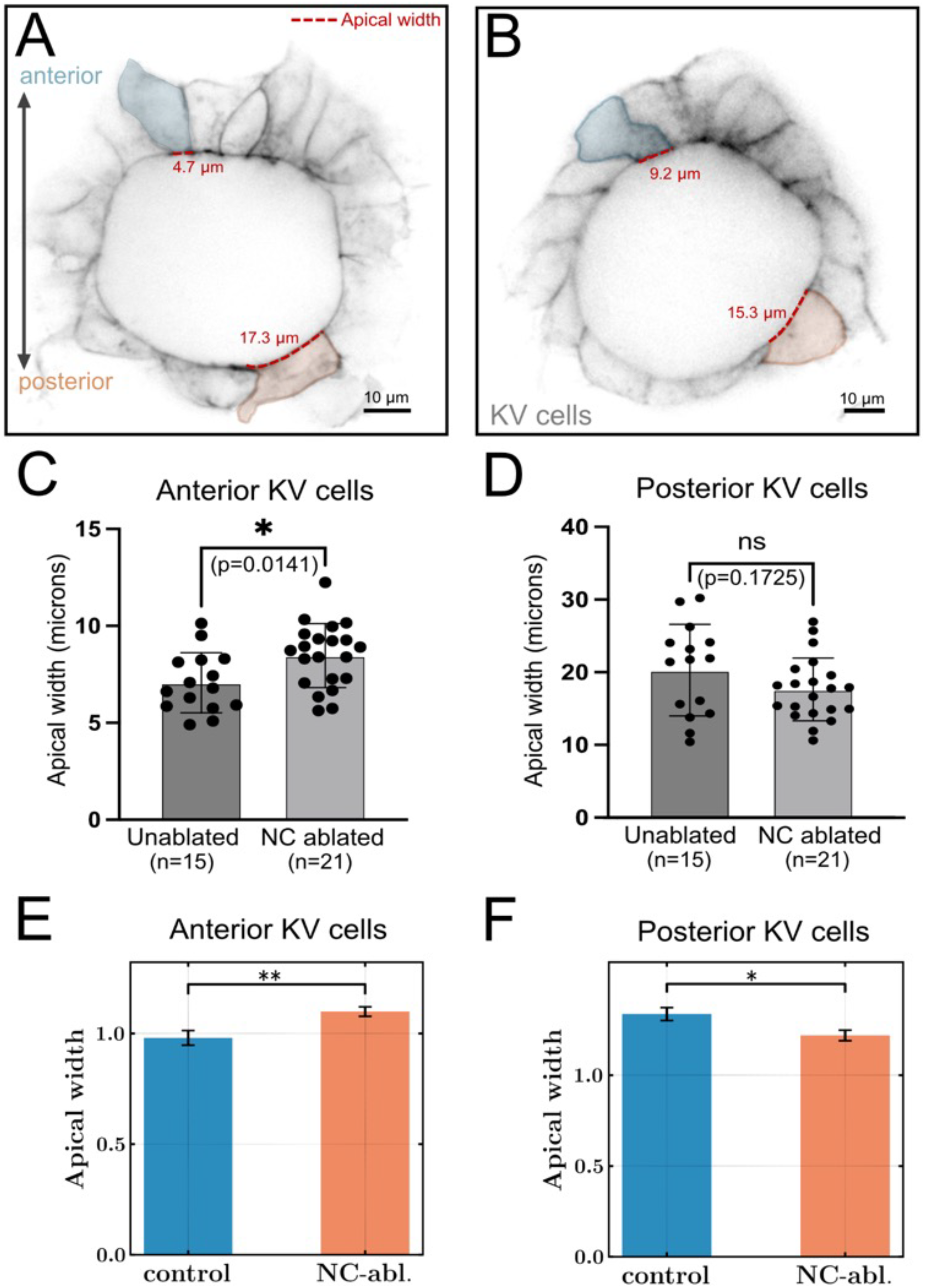
KV cell shapes are altered by notochord ablation. (A-B) Confocal microscopy images of the midplane of KV with GFP-labeled KV cell membranes (gray) in a live unablated embryo (A) and a notochord (NC) ablated embryo (B) at the 8 somite stage. Representative anterior (blue) and posterior (orange) KV cells are highlighted with measurements of apical cell width (dashed red line). (C-D) The average apical width of anterior KV cells (C) and posterior KV cells (D) in control and NC ablated embryos. Each point represents an embryo. 70 anterior KV cells and 39 posterior KV cells were measured in 15 unablated control embryos and 100 anterior KV cells and 61 posterior KV cells were measured in 21 notochord ablated embryos that were pooled from 3 independent experiments. Error bars=1 standard deviation. ns=not significant. *p<0.05. P values are indicated on each graph. (E-F) Average apical width (in natural simulation units) of anterior KV cells (E) and posterior KV cells (F) from 3D vertex simulations for control and NC ablation conditions. As in the experiments, apical width was measured from the midplane of the KV at steady state across 17 independent simulation runs. A total of 125 anterior and 98 posterior KV cells were analyzed in control simulations, and 116 anterior and 104 posterior KV cells in NC ablation simulations.

## Discussion and conclusions

We have demonstrated that zebrafish Kupffer’s vesicle, a useful model for probing mechanisms of organ morphogenesis, develops a stereotyped oblate shape in wildtype embryos, and simultaneously moves dynamically through the tissue with a stereotyped variation in speed. A 3D mechanical vertex model predicts that self-generated tissue-scale dynamic forces associated with this motion of KV generate the observed organ shape, as well as individual cell shape changes. The model further predicts how cell and organ shapes will quantitatively change if the dynamical forces are altered. We tested this hypothesis with laser ablation experiments that perturb those dynamical forces and found altered organ shapes and cilia densities that were consistent with model predictions. Independent of specific model predictions, our experiments demonstrate that tissue-scale dynamic forces help govern cell and organ shape changes.

This is a surprising result. Although there are many studies that have used hydrodynamic models with drag forces to explain large-scale tissue flows and shapes ^12, 13, 42, 43^, much previous work on cell-based mechanical models(including our own) has assumed that motion in development is slow enough that we can consider tissues to be roughly in force balance (e.g. unbalanced forces are small and negligible in determining tissue morphology)^2, 4, 6, 7, 9^. While a few recent studies have included effects of frictional forces as boundary conditions on cell-based models ^10, 11^, these studies have remained somewhat disconnected from the hydrodynamic models that emphasize the role of bulk tissue flows. Our work here confirms that small velocities in bulk tissue-scale flow can generate forces that are significant enough to locally deform cells and organs, ultimately controlling their shape. This makes sense because the timescale of tissue relaxation is large in the zebrafish tailbud, since it is near a fluid-solid transition.^28^ Given that recent work indicates fluid-solid transitions seem to occur quite frequently in development,^3, 4^ our results highlight that self-generated bulk viscous forces should not be ignored, and may be playing a role in many other developmental processes.

It is important to note that while we have demonstrated that dynamical forces play a role in driving asymmetric AP cilia distribution, and therefore the remodeling of individual KV cell shapes, these dynamical forces are clearly not the only mechanism driving those processes. KV remodeling still occurs (albeit less strongly), and ultimately downstream LR patterning is still successful in ablated embryos. This suggests multiple mechanisms (perhaps including cell interactions with ECM at the interface with the notochord^27^) are acting to robustly drive this process.

A common theme in development is that multiple partially redundant mechanisms act to drive morphogenetic processes that are important for fitness^44-46^. Our combination of experiments and modeling provides a framework for teasing apart the contributions of such redundant mechanisms. Future work could extend this framework to knock down multiple mechanisms simultaneously and predict how that may ultimately abrogate LR patterning. More broadly, this combination modeling/experiment approach could be used to distinguish multiple mechanisms in other developmental systems.

## Acknowledgements

We thank Elizabeth Lawson-Keister and Sadjad Arzash for helpful discussions, and Tao Zhang and Jen Schwarz for 3D vertex model code. This work was primarily supported by NIH R01HD099031. We also acknowledge R01GM-127621 (H.H.) and no. R01GM-130874 (H.H.), and NSF-CMMI-1334611 (M.L.M).

## Author contributions

Conceptualization: RKM, MLM, JDA; Methodology: RKM, MLM, EMR, MB, YL, AMH, HH, JDA, OAR; Investigation: RKM, EMR, MB, AMH, OAR; Visualization: RKM, EMR, AMH; Funding acquisition: MLM, JDA, HH; Project administration: MLM, JDA; Supervision: MLM, JDA, HH; Writing – original draft: RKM, MLM, EMR, JDA; Writing – review & editing: RKM, MLM, EMR, JDA, AMH, YL, OAR, HH.

## Data and materials availability

The data and code necessary to reproduce the analyses of this manuscript are available at Dryad.

## Supplementary Materials

Materials and Methods

Figs. S1 to S22

Movies S1 to S11

Tables S1 to S7

References*(1–12)*

