## Supplemental Text and Figures for "Dynamical forces drive cell and organ morphology changes during embryonic development"

**The PDF file includes:**

Materials and Methods
Figs. S1 to S18
Tables S1 to S7
References

**Other supporting materials for this manuscript include the following:**

Movies S1 to S11

### Materials and Methods

#### *Zebrafish strains and husbandry*

Transgenic zebrafish (*Danio rerio*) strains used in this study are: *Tg(ubb:LOXP-mScarlet-NLS-LOXP-QF-GAL4)$^{hsc150}$* (*1*) obtained from Dr. Bret Pearson, *Tg(-2.4shha-ABC:GFP)*$^{sb15}$ (*2*) obtained from ZIRC, and *Tg(sox17:EGFP-CAAX,myl7:EGFP)$^{sny101}$* that we generated previously (*3*). Adult zebrafish were maintained in an aquarium rack system on a 14 hour light/10 hour dark cycle. Embryos were collected from natural matings and staged as described (*4*). All experiments using zebrafish were approved by the International Animal Care and Use Committee at SUNY Upstate Medical University.

#### *Time-lapse imaging of zebrafish embryos*

Zebrafish embryos were incubated at 28.5°C until they reached the tailbud stage of development. Embryos were then dechorionated and immobilized in 1.5% low melting point agarose in glass bottomed culture dishes (MatTek). Immobilized embryos were imaged between the 2 and 8 somite stages on a Leica Deconvolution system (DMi8) with X-LightV2 spinning disk confocal microscope using a 40X objective. A Z-series of images (2 µm steps) through KV in the tailbud (dorsal view) was captured every 2 minutes for up to 2 hours of development. Under these conditions, embryos develop a new somite pair every 30 minutes. After imaging, embryos were removed from the agarose and allowed to develop for downstream analyses. 3D reconstructions of the images were visualized using Imaris software (Bitplane). To create time-lapse movies, ImageJ software (NIH) was used to make maximum Z projections using 40 µm above and below the KV for a total depth of 80 µm through the tailbud.

#### *Laser ablation of cells in zebrafish embryos*

To ablate specific cells in a zebrafish embryo, we used an X-light v2 Confocal Unit spinning disk confocal microscope with VisiView kinetics unit coupled to a 355 nm pulsed laser with a 40X 1.15 NA water objective. For notochord cell ablations, we used transgenic *Tg(sox17:EGFP-CAAX,myl7:EGFP),Tg(ubb:LOXP-mScarlet-NLS-LOXP-QF-GAL4)*, and *Tg(-2.4shha-ABC:GFP)* embryos that express membrane-localized EGFP in KV cells, nuclear-localized mScarlet in all cells, and cytoplasmic GFP in notochord cells, respectively At the tailbud stage, embryos were dechorionated and immobilized in glass bottomed dishes (MatTek). Between the 2 to 3 somite stages, a rectangular region of interest (ROI) measuring approximately 85 $\mu m$ in length was used to ablate a section of notochord cells positioned approximately 5-10 cell diameters away from KV. To ablate posterior KV cells, we used double transgenic *Tg(sox17:EGFP-CAAX,myl7:EGFP)*; *Tg(ubb:LOXP-mScarlet-NLS-LOXP-QF-GAL4)* embryos that allow visualization of KV cell membranes in the green channel and nuclei in the red channel. A custom drawn ROI was used to ablate 3 to 8 cells in the posterior region of the KV at the middle focal plane (defined as the plane with the largest lumen diameter) between the 2 to 3 somite stages. For control ablations, an ROI was used to ablate 3 to 8 cells in the anterior head region of the embryo far away from KV, notochord, and tailbud. To visualize movement of KV through the tailbud following cell ablation, time-lapse imaging was conducted as described above.

#### *KV lumen shape calculation*

3D reconstructions of confocal microscopy time-lapse images of KV were generated using Imaris software (Bitplane). These reconstructions were used to uniformly align KV with respect to anterior-posterior and left-right axes. The OrthoSlicer tool in Imaris was then used to isolate the middle plane of KV, which we define as the plane with the largest lumen diameter. The outline of the KV lumen shape in the middle plane was traced using Fiji software (NIH). XY coordinates for this shape were then used in custom Python codes to obtain the radius of gyration ($R_g$) of the lumen. The radius of gyration of the lumen is defined as

$$Rg = \left[\frac{1}{N}\sum_{i=1}^{N} |\boldsymbol{r}_i - \boldsymbol{r}_{cm}|^2\right]^{1/2} .$$

Here, $N$ represents the number of boundary points of the lumen, $r_i(x_i, y_i)$ denotes the position of each boundary point and $\boldsymbol{r}_{cm} = \frac{1}{N}\sum_{i=1}^{N} \boldsymbol{r}_i$ represents the center of mass position of the lumen. We take y-axis along the anterior-posterior (AP) axis and left-right (LR) axis is along x-axis. The radius of gyration along AP-axis is similarly defined as

$$Rg_{AP} = \left[\frac{1}{N}\sum_{i=1}^{N} |y_i - y_{cm}|^2\right]^{1/2} ,$$

where $y_i$ and $y_{cm}$ are the position of $i^{th}$ point and center of mass of the lumen along the y-axis respectively. Similarly, the radius of gyration along x-axis is defined using x-component of position vectors. We calculate a shape parameter ($Sr$) defined as the ratio between the radius of gyration of the lumen along the AP-axis ($Rg_{AP}$) and along the LR-axis ($Rg_{LR}$),

$$Sr = \frac{Rg_{AP}}{Rg_{LR}} .$$

This shape parameter, $Sr$, is utilized to determine the lumen's shape changes in comparison to a perfect sphere, where the value of $Sr$ would be 1. Data from three time points (separated by 2 mins) were averaged for each somite stage.

***KV lumen speed calculation***

To calculate the velocity of KV moving through the tailbud, the outline of the KV lumen shape in the middle plane was traced using Fiji software (NIH). A custom Python code was used to identify the center of mass of the KV lumen. One center of mass calculation was made for KV at each somite stage during time-lapse experiments. The velocity of KV lumen at somite stage, $s = n + 1/2$ is then determined by computing distance travelled by lumen between two consecutive somite stages,

$$u\left(s = n + \frac{1}{2}\right) = \frac{|r_{cm}(s = n + 1) - r_{cm}(s = n)|}{\Delta t} .$$

Here $\Delta t$ represents the time between two somite stages (which is 30 minutes) and $r_{cm}(s = i)$ denotes the center of mass position of the lumen at the $i^{th}$ somite stage. The velocity is expressed in micrometers per minute (*μm/min*).

***Fluorescent immunostaining, RNA in situ hybridization, and scoring heart laterality***

For immunostaining of KV cilia, embryos were fixed overnight at 4° C in 4% paraformaldehyde in PBS with 0.5% Triton X-100 and then processed for whole mount antibody staining as described (*5*). Primary antibodies used were anti-EGFP (GeneTex, GTX13970) diluted to 1:400

to detect KV cells in *Tg(sox17:EGFP-CAAX,myl7:EGFP)* embryos and anti-acetylated tubulin (Sigma T6793) diluted 1:200 to label cilia. Alexa Fluor 488 or 568 fluorescent secondary antibodies (Invitrogen) were used at 1:200 dilutions. Embryos were imaged using a Nikon spinning disk confocal microscope system (Yokogawa CSU-X1) using a 40x objective. Individual KV cilia were identified and marked in blinded 3D images using Imaris software (Bitplane). When necessary, images were rotated for uniform orientations with anterior at top. The number of ciliated cells located in the anterior and posterior regions of KV (as defined by a line that bisects KV along the AP axis at the widest lumen diameter) was determined. A ratio of the number of anterior ciliated cells to the number of posterior ciliated cells (Na/Np) was calculated for each embryo. An unpaired two-tailed t-test with Welch's correction was used for statistical analysis. A P value less than 0.05 was considered a significant difference. Whole-embryo RNA *in situ* hybridization analysis of southpaw (*spaw)* expression in lateral plate mesoderm was performed as previously described (*6*). Heart looping was visualized in live embryos at 2 days post-fertilization under a dissecting microscope. Hearts were labeled with EGFP from myl:EGFP transgene expression.

***Analysis of KV cell shapes in experiments and simulations***

Images of labeled KV cells in *Tg(sox17:EGFP-CAAX,myl7:EGFP)* embryos were captured at the 8-somite stage using a Nikon spinning disk microscope. Blinded images were processed using Imaris software. The oblique slicer tool was used to isolate the midplane of KV (defined as the plane in which the lumen has the largest diameter). The midplane was divided into 4 equal quadrants: Anterior, Posterior, left, and Right. Cells in the Anterior and Posterior quadrants were analyzed. The apical width (apical surface along the lumen) and height of the cell were measured using the segmented line tool in ImageJ software. If either the width or height of a cell could not be resolved, the cell was excluded from the analysis. An unpaired two-tailed t-test with Welch's correction was used for statistical analysis. A P value less than 0.05 was considered a significant difference.

Cell shapes in simulations were measured using techniques that were as similar as possible to those used in experiments. The apical width of KV cells in 3D vertex simulations were measured at the midplane of the KV. Cell shapes at the midplane were extracted using a slice filter in ParaView software. The midplane was divided into anterior and posterior regions based on a horizontal line passing through the center of the KV. Cells intersecting this horizontal line were excluded from the analysis. The apical width of KV cells along the lumen interface was measured using ImageJ software using the same methods as described for the experiments in the previous paragraph.

***3D Vertex model for KV motion in tailbud tissue***

We utilize a 3D Vertex model (*7*, *8*) to capture the propulsion of KV through tailbud tissue. The KV is modeled as one layer of epithelial cells distributed over a spherical surface that encloses the lumen (Fig. 1D-E). In the zebrafish embryo, the lumen is a fluid-filled membrane, and in our model, we represent that roughly spherical volume with $N_L = 17$ model cells (marked by yellow). The KV-lumen system is embedded in the three-dimensional cubic domain of tailbud cells ($N_T = 2048$, marked by grey). KV is composed of $N_{KV} = 50$ cells (marked by green) and anterior and posterior parts of the lumen are marked by orange and purple, respectively. In our model, each cell is represented by a deformable polyhedron, and the tissue comprises the space filled by such polyhedrons. Unlike the Voronoi model (*9*, *10*), where the cell centers are the degrees of freedom, in vertex models, the degrees of freedom are the cell vertices. This additional degree of freedom is crucial for more accurately capturing realistic shape changes in systems with heterogeneous architectures like the KV in tailbud tissue (*11*). The energy of a confluent tissue in a 3D vertex model with $N$ cells is given by

$$E = \sum_{i=1}^{N} \left[ K_V (V^i - V_0^i)^2 + K_S (S^i - S_0^i)^2 \right] + \sum_{i \neq j} \gamma^{ij} A^{ij} \,,$$

where the first term ensures the volume incompressibility of the cell, and $V^i$ and $V_0^i$ are the actual and preferred volume of the $i^{\text{th}}$ cell, respectively. The second term penalizes departures of the actual surface area ($S^i$) of the $i^{\text{th}}$ cell from its preferred surface area $S_0^i$. This surface energy term represents effective mechanical interaction due to the competition between cell-cell adhesion and actomyosin contractility in the cytoskeleton localized near the cell cortex. $K_V$ and $K_S$ are respectively the moduli that penalize changes to the volume and surface area. The last term captures additional heterotypic surface tension, $\gamma^{ij}$, between cell i and cell j of different cell types that share a surface area $A^{ij}$. This heterotypic surface tension between two different tissues ensures a mechanical boundary between two different tissue types that is necessary to capture experimental observations of cell sorting in heterotypic mixtures (*11*, *12*). In our model, it creates boundary between lumen and KV and KV and tailbud tissue.

The mechanical force on each vertex is then obtained from the gradient of the energy $E$, $\boldsymbol{F}_i^c = -\nabla_i E$. The anterior and posterior part of the KV is also subjected to additional dynamic (active) forces, $\boldsymbol{F}_i^d$ due to pushing forces from notochord and pulling forces from posterior part of the KV. The motion of each cell is governed by an over-damped equation,

$$\frac{d\boldsymbol{r}_i}{dt} = \mu \boldsymbol{F}_i^c + \mu \boldsymbol{F}_i^d + \sqrt{2\mu k_B T}\boldsymbol{\zeta}_i.$$

Here $\zeta$ is the spatially isotropic Gaussian white noise with unit variance, such that in the absence of dynamic forces, the system obeys the fluctuation-dissipation theorem. $\mu$ is the mobility and $T$ is the effective temperature of the system. In the absence of the activity the diffusion coefficient of each vertex is $\mu k_B T$.

We apply the dynamic (active) forces only on vertices that are part of the anterior and posterior part of KV. The vertices of the anterior part of the KV (marked by orange color in Fig. 1D-E of the main text) is determined by the distance (*r*) of the vertices from AP axis through center of the lumen. If the distance of the vertices (*r*) is less than the notochord radius ($r_{nc} = 1$), we take those vertices as a part of the anterior part of the KV. The vertices on the posterior part of the KV are chosen such that azimuthal angle for the vertices, $\varphi_{pc} = cos^{-1}(\Delta y / r_1)$ is greater than $0.75\pi$. Here $\Delta y$ is the distance of the vertices along y-axis from the center of the lumen and $r_1$ is absolute distance of the vertices from center of the lumen. The total dynamic forces on the anterior and posterior part of the KV is given by respectively $-f_A \hat{\boldsymbol{e}}_y$ and $-f_P \hat{\boldsymbol{e}}_y$. We update the number of vertices on the anterior ($v_A$) and posterior ($v_P$) part of the cells at each simulation time. The amount of external forces on the posterior (anterior) part of the KV are equally distributed on the all vertices in the posterior (anterior) part. With these specifications, the dynamic forces on the anterior and posterior part of KV is then given by

$$\boldsymbol{F}_{\boldsymbol{i}}^{\boldsymbol{d}} = \begin{array}{l} -(f_A/v_A)\hat{\boldsymbol{e}}_y \ \ anterior, \\ -(f_P/v_P)\hat{\boldsymbol{e}}_y \ \ posterior. \end{array}$$

We take the target volume $V_0^i$ to be 1 for all cells. The target surface area for KV and lumen cells are taken as $S_0 = 5.0$ and we vary the target surface area of tailbud cells. We have used two different simulation protocols, one to initialize the system and a second to simulate its dynamics. The initialization of KV-lumen-tailbud cells system is obtained from the following steps: a) We first use random Voronoi tessellation points to get cells vertices and polygons with uniform mechanical properties, b) then we choose 17 cells at the center of the domain to represent lumen cells, c) the KV cells are then assigned at the boundary of the lumen cells such that total number of KV cells are 50. We first equilibrate KV-lumen-tailbud cells without active forces such that system reaches a minimum energy configuration. Next, we run the simulation with the active forces. We use Euler scheme to integrate the equation of motion with time steps $5 \times 10^{-2}$(for equilibration) and $10^{-3}$ (with active forces). Each simulation is run for $6.1 \times 10^6$ time steps. The simulation data in Fig 2I-J, Fig 3E and Fig S15 are obtained by averaging 18 independent runs. The energy, time and length are nondimensionalized respectively by natural units $K_S {V_0}^{4/3}$, $1/(\mu K_S {V_0}^{2/3})$, and ${V_0}^{1/3}$. All simulation results are presented in these natural units. We take $\mu = 1$ and $k_B T = 10^{-5}$. The parameters for the simulations are given in Table S4-S6.

***Hydrodynamic Model for Lumen Shape Change***

To understand whether the influence of individual forces on the shape alterations of the lumen depends on specific features of the vertex model or is a generic consequence of those active forces, we devised a hydrodynamic model of lumen motion and shape. The middle plane of lumen is considered as a deformable two-dimensional membrane composed of $N$ connecting spherical beads of radius $b$ and position $r_i$. The body force $F_i^b$ acting on the i[th] bead is obtained from the gradient of total potential U,

$$F_i^b = -\nabla U.$$

Here, the total potential is the sum of connectivity ($U^C$), bending ($U^B$), self-avoidance ($U^S$) and area potential ($U^A$),

$$U = \sum_{i=1}^{N} U^C(\boldsymbol{r}_i, \boldsymbol{r}_{i+1}) + \sum_{i=1}^{N} U^{BC}(\boldsymbol{r}_{i-1}, \boldsymbol{r}_i, \boldsymbol{r}_{i+1}) + \sum_{i<j}^{N} U^S(\boldsymbol{r}_i, \boldsymbol{r}_j) + \sum_{i=1}^{N} U^A(\boldsymbol{r}_1, .., \boldsymbol{r}_N),$$

where $\boldsymbol{r}_{N+1=} \boldsymbol{r}_1$ and $\boldsymbol{r}_{-1=} \boldsymbol{r}_N$. The two body connectivity potential is harmonic spring potential $U^C = k(r - b_0)^2$ where $b_0 = 2b$ is the equilibrium bond length and $r = |\boldsymbol{r}_{i+1} - \boldsymbol{r}_i|$. The three-body bending potential is $U^B = \kappa(1 - \cos(\phi))$, which penalizes departures of the angle ($\phi$) between consecutive bond vectors from its equilibrium value of zero. Here, $\kappa$ is the bending stiffness of the membrane. The self-avoidance potential between beads is considered as Weeks-Chandler-Anderson potential of strength $\epsilon$ which vanishes if the distance between beads $r_{ij} = |\boldsymbol{r}_i - \boldsymbol{r}_j|$ exceeds 2b. To keep the area ($A$) of the lumen conserved, we use area potentials $U^A = K_A(A - A_0)^2$, where $K_A$ is area-stretching modulus and $A_0$ is the preferred area of the membrane.

In addition to the body force, the beads at the anterior (marked by orange in Fig. S12A) and posterior part of the membrane (marked by purple in Fig. S12A) is subjected to external forces, $\boldsymbol{F}_i^e = -f\ \hat{e}_{yz}$.

With these descriptions of the forces, the dynamics of each bead in the membrane can be described as overdamped equation of motion (*13*)

$$\dot{\boldsymbol{r}}_i = \boldsymbol{\mu}_{\boldsymbol{ij}} \cdot (\boldsymbol{F}_j^b + \boldsymbol{F}_j^e),$$

where $\boldsymbol{\mu}_{\boldsymbol{ij}}$ is the mobility tensor,

$$8\pi\eta\boldsymbol{\mu}_{\boldsymbol{ij}}(\boldsymbol{r}_i, \boldsymbol{r}_j) = \begin{cases} \left(1 + \frac{b^2}{6}\boldsymbol{\nabla}_i^2\right)\left(1 + \frac{b^2}{6}\boldsymbol{\nabla}_j^2\right)\boldsymbol{G}(\boldsymbol{r}_i, \boldsymbol{r}_j), & i \neq j, \\ \frac{4}{3b}\boldsymbol{\delta}_{ij}, & i = j. \end{cases}$$

Here $\boldsymbol{G}(\boldsymbol{r}_i, \boldsymbol{r}_j)$ is Green's function for Stokes flow and $\eta$ is the viscosity of the medium.

$$\boldsymbol{G}_{\boldsymbol{ij}} = \frac{\boldsymbol{\delta}_{ij}}{\rho} + \frac{\boldsymbol{\rho}_i\,\boldsymbol{\rho}_j}{\rho},$$

with $\boldsymbol{\rho} = \boldsymbol{r}_i - \boldsymbol{r}_j$ and $\rho = |\boldsymbol{\rho}|$. The magnitude of external forces ($f$) on the anterior and posterior part of the KV are respectively $f_A$ and $f_P$. Similar to the 3D vertex model, at each simulation time step we identify beads that are part of the anterior and posterior part of the membrane. The magnitude of external forces on the posterior (anterior) part of the membrane are equally distributed on all beads in the posterior (anterior) part. The equation of motion is integrated by Euler method with time step $10^{-2}$. The simulation parameters are given in the Table S7.

***Data availability***

Data associated with this manuscript will be made available as a Dryad repository.

### Supplemental Figures

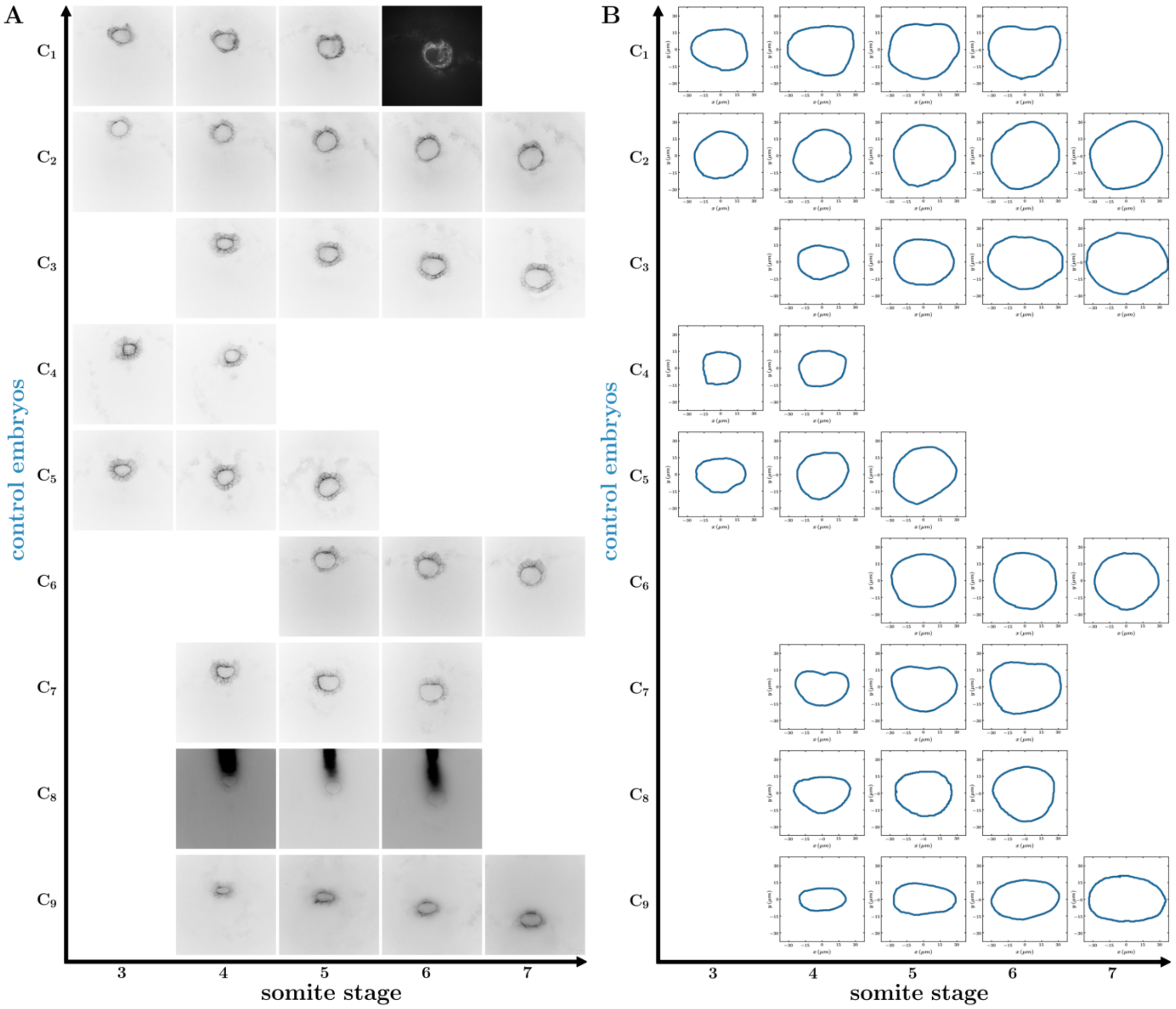


**Fig. S1. KV middle plane and the corresponding lumen boundary for the computation of KV speed in unablated control experiments.** (A) The motion of the KV in the tailbud tissue (shown by the middle plane of KV) as a function of somite stage for different unablated control embryos (marked $C_1$ to $C_9$). (B) The boundary of the lumen obtained from the middle plane of the KV is depicted in the right-side panel.

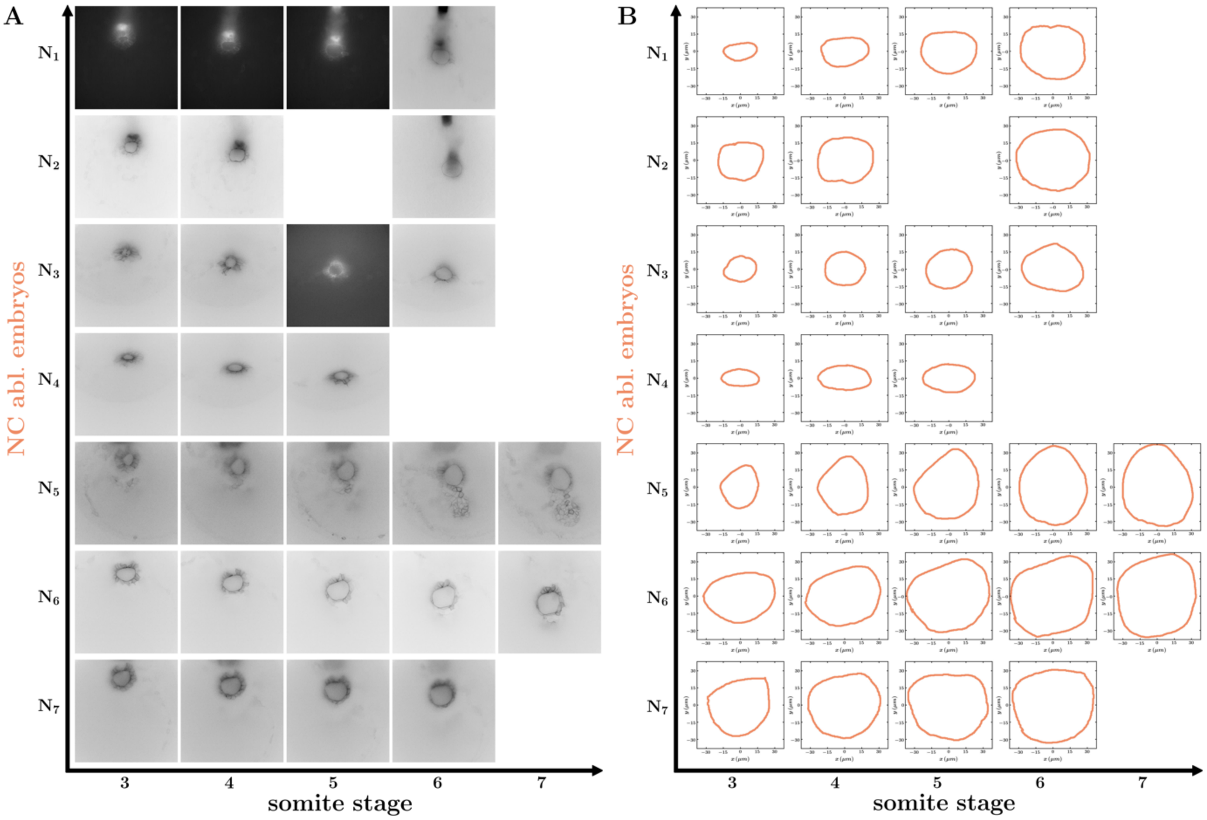


**Fig. S2. KV middle plane and the corresponding lumen boundary for the computation of KV speed in notochord ablation experiments.** (A) The motion of the KV in the tailbud tissue (shown by the middle plane of KV) as a function of somite stage for different notochord ablation embryos (marked $N_1$ to $N_7$). (B) The boundary of the lumen obtained from the middle plane of the KV is depicted in the right-side panel.

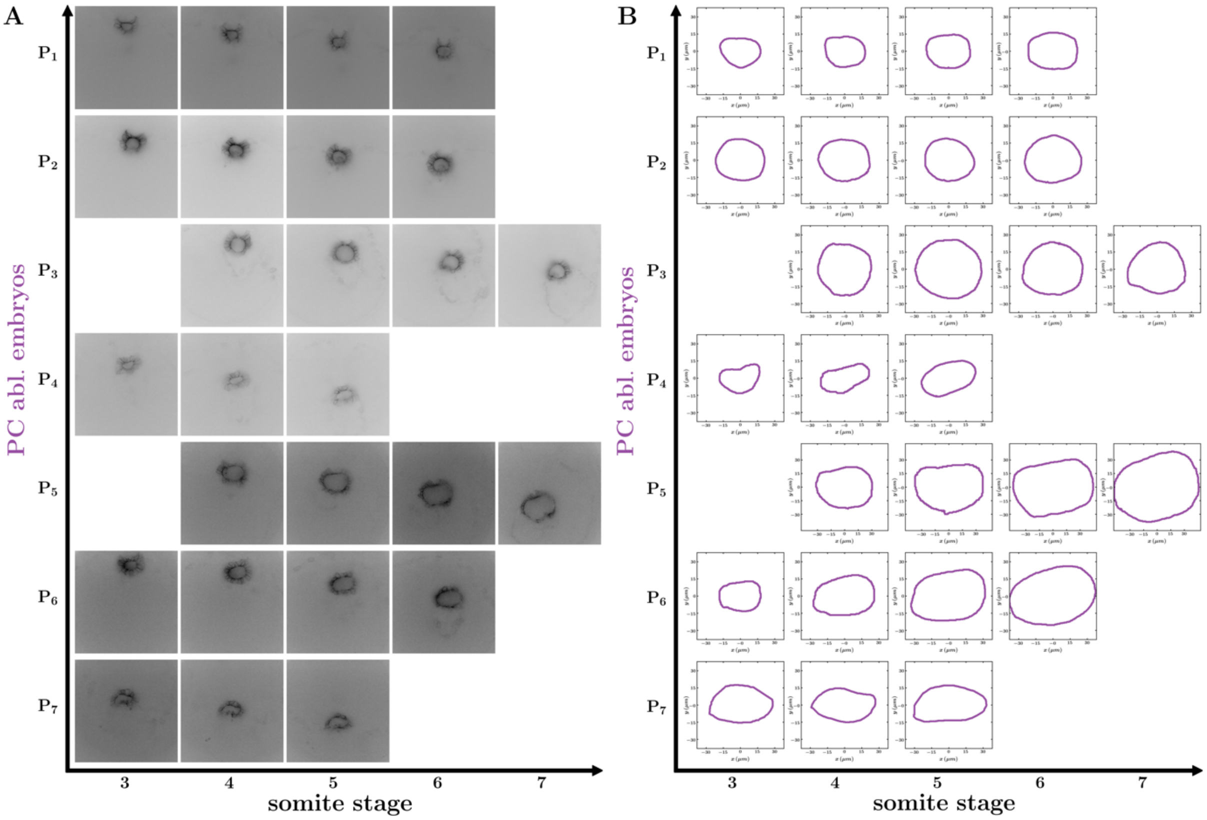


**Fig. S3. KV middle plane and the corresponding lumen boundary for the computation of KV speed in posterior cells ablation experiments.** (A) The motion of the KV in the tailbud tissue (shown by the middle plane of KV) as a function of somite stage for different posterior cells ablation embryos (marked $P_1$ to $P_7$). (B) The boundary of the lumen obtained from the middle plane of the KV is depicted in the right-side panel.

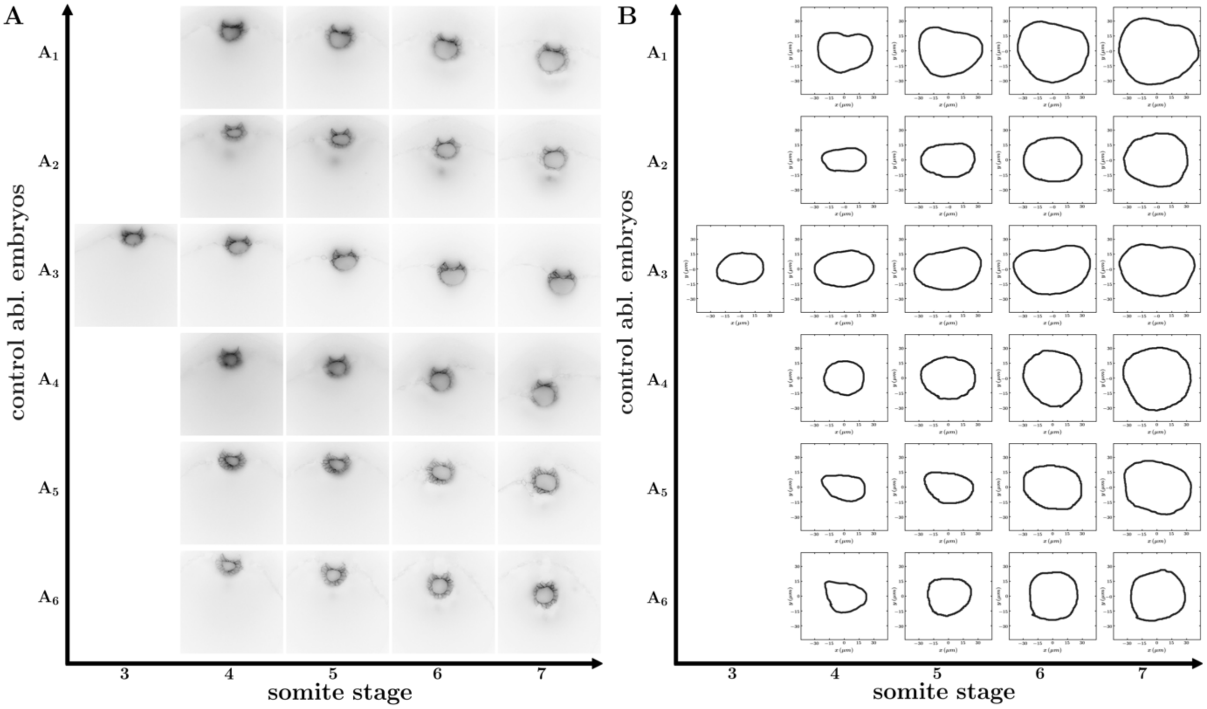


**Fig. S4. KV middle plane and the corresponding lumen boundary for the computation of KV speed in control ablation experiments.** (A) The motion of the KV in the tailbud tissue (shown by the middle plane of KV) as a function of somite stage for different posterior cells ablation embryos (marked $P_1$ to $P_7$). (B) The boundary of the lumen obtained from the middle plane of the KV is depicted in the right-side panel.

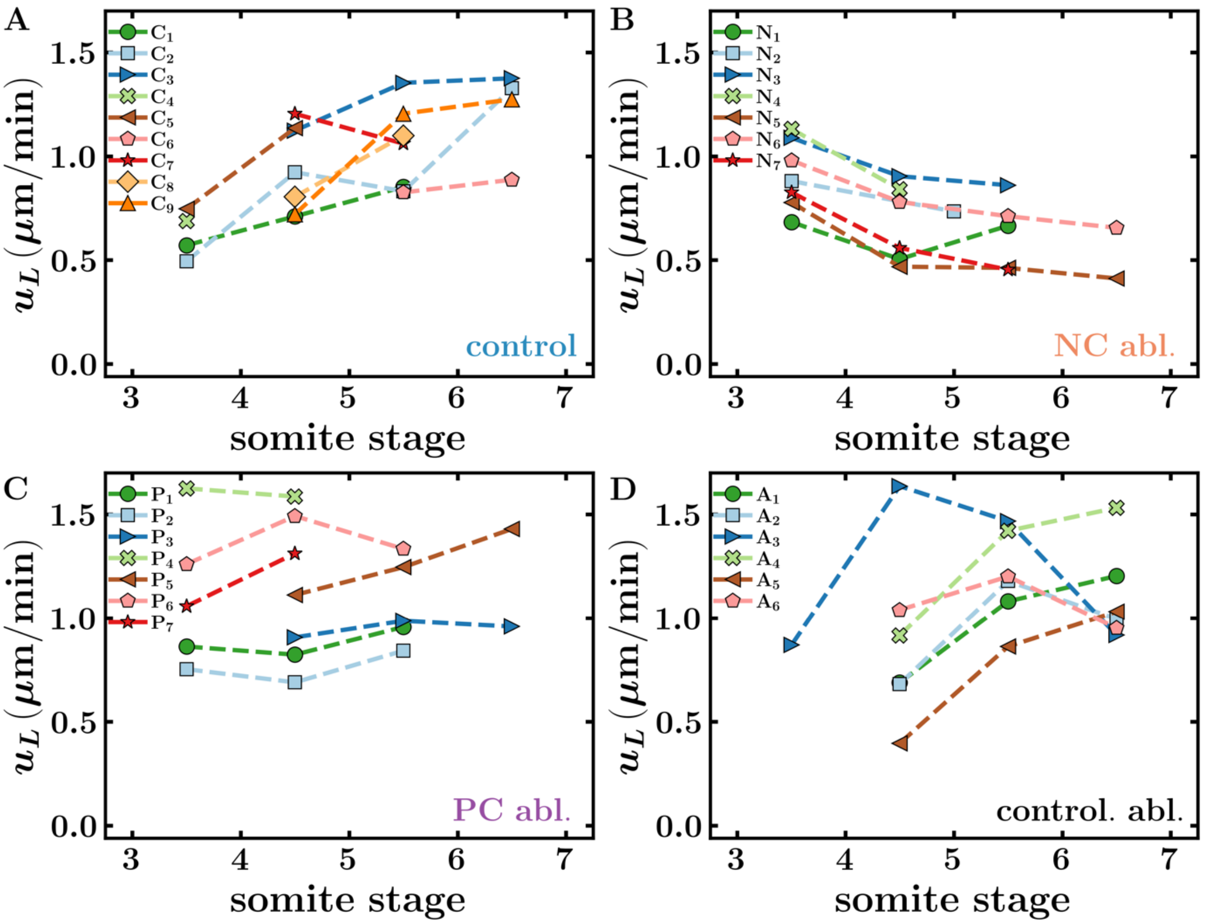


**Fig. S5. The speed of KV lumen in control and different ablation experiments.** The speed of individual KV lumen as a function of somite stage in unablated control (A), notochord ablation (B), posterior cells ablation (C), and control ablation (D) experiments.

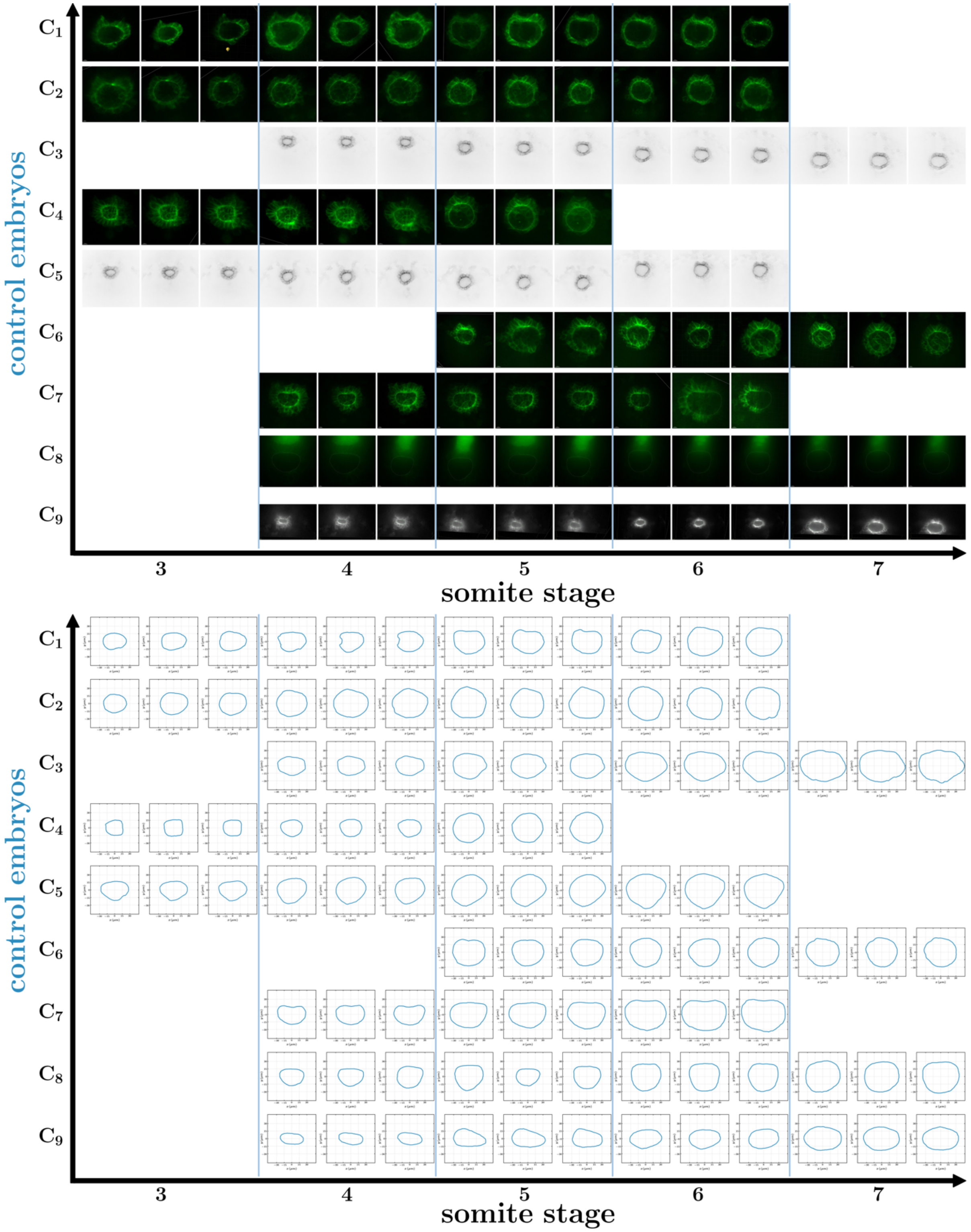


**Fig. S6. KV middle plane (A) and the corresponding lumen boundary (B) for the computation of lumen shape in unablated control experiments.** Different embryos are marked by $C_1$ to $C_9$. To calculate the lumen shape at each somite stage, three time snapshots separated by 2 minutes (0.033 somite stage) are used.

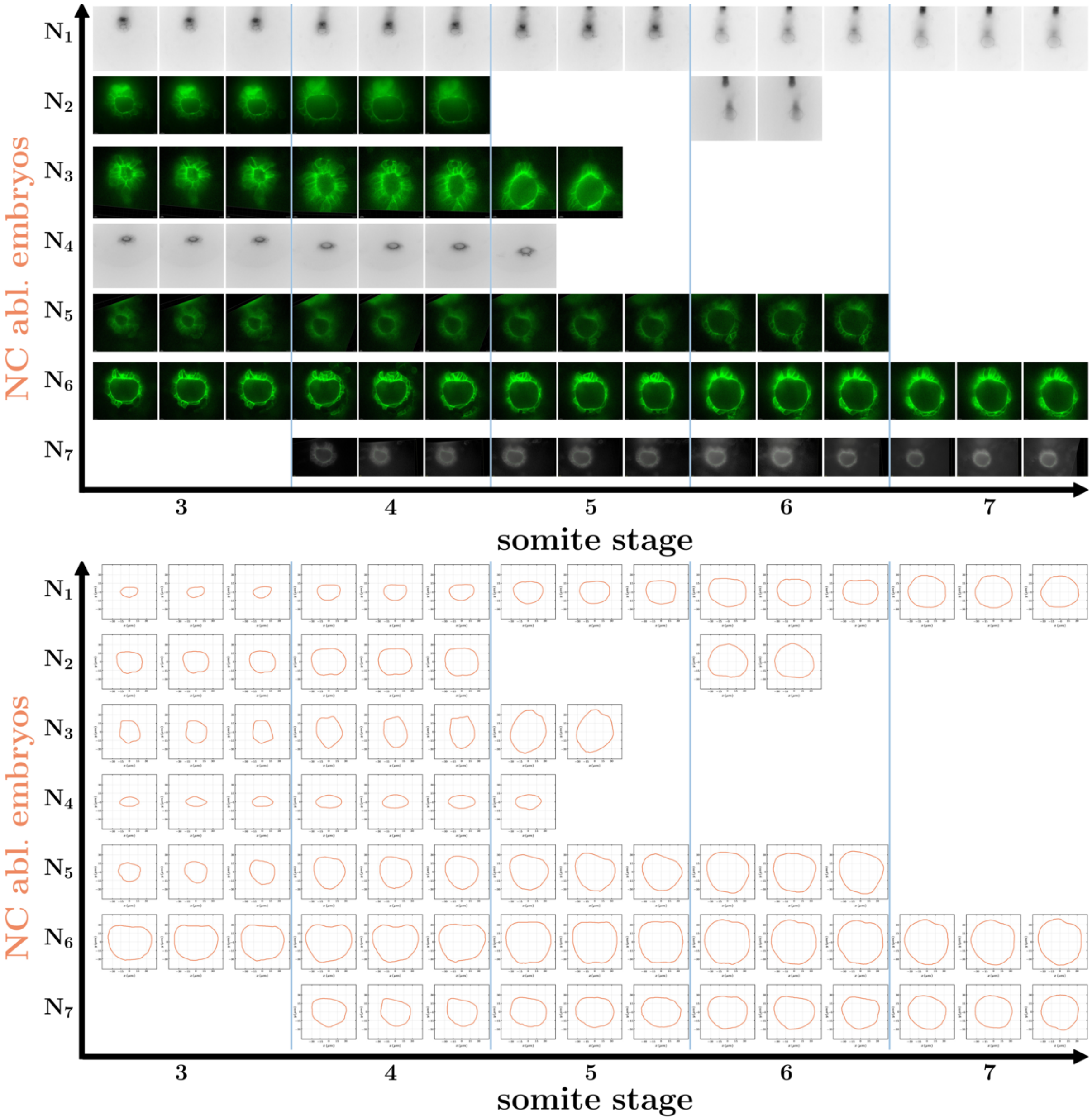


**Fig. S7. KV middle plane (A) and the corresponding lumen boundary (B) for the computation of lumen shape in notochord ablation experiments.** Different embryos are marked by $N_1$ to $N_7$. To calculate the lumen shape at each somite stage, three time snapshots separated by 2 minutes (0.033 somite stage) are used.

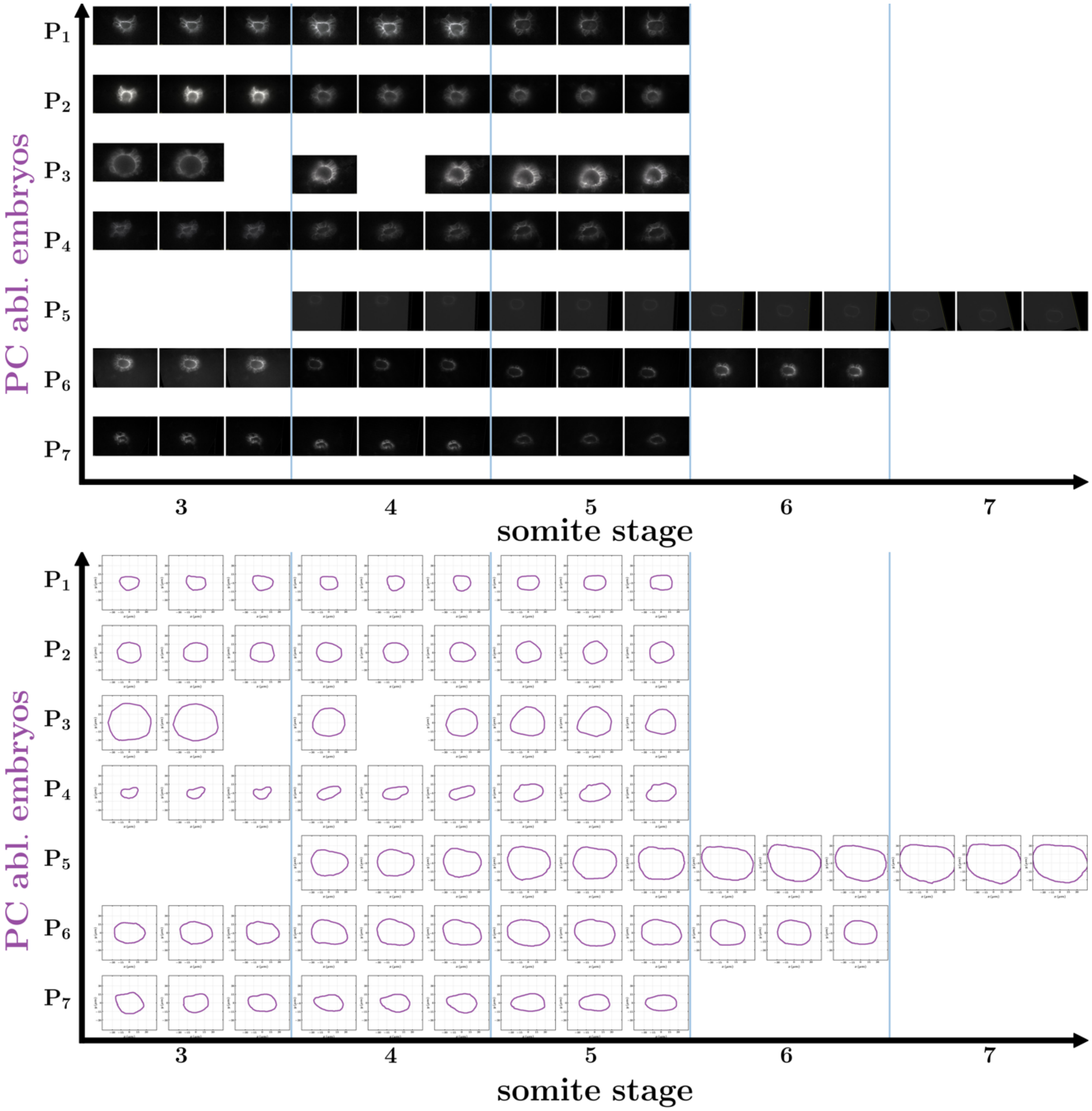


**Fig. S8. KV middle plane (A) and the corresponding lumen boundary (B) for the computation of lumen shape in posterior cells ablation experiments.** Different embryos are marked by $N_1$ to $N_7$. To calculate the lumen shape at each somite stage, three time snapshots separated by 2 minutes (0.033 somite stage) are used.

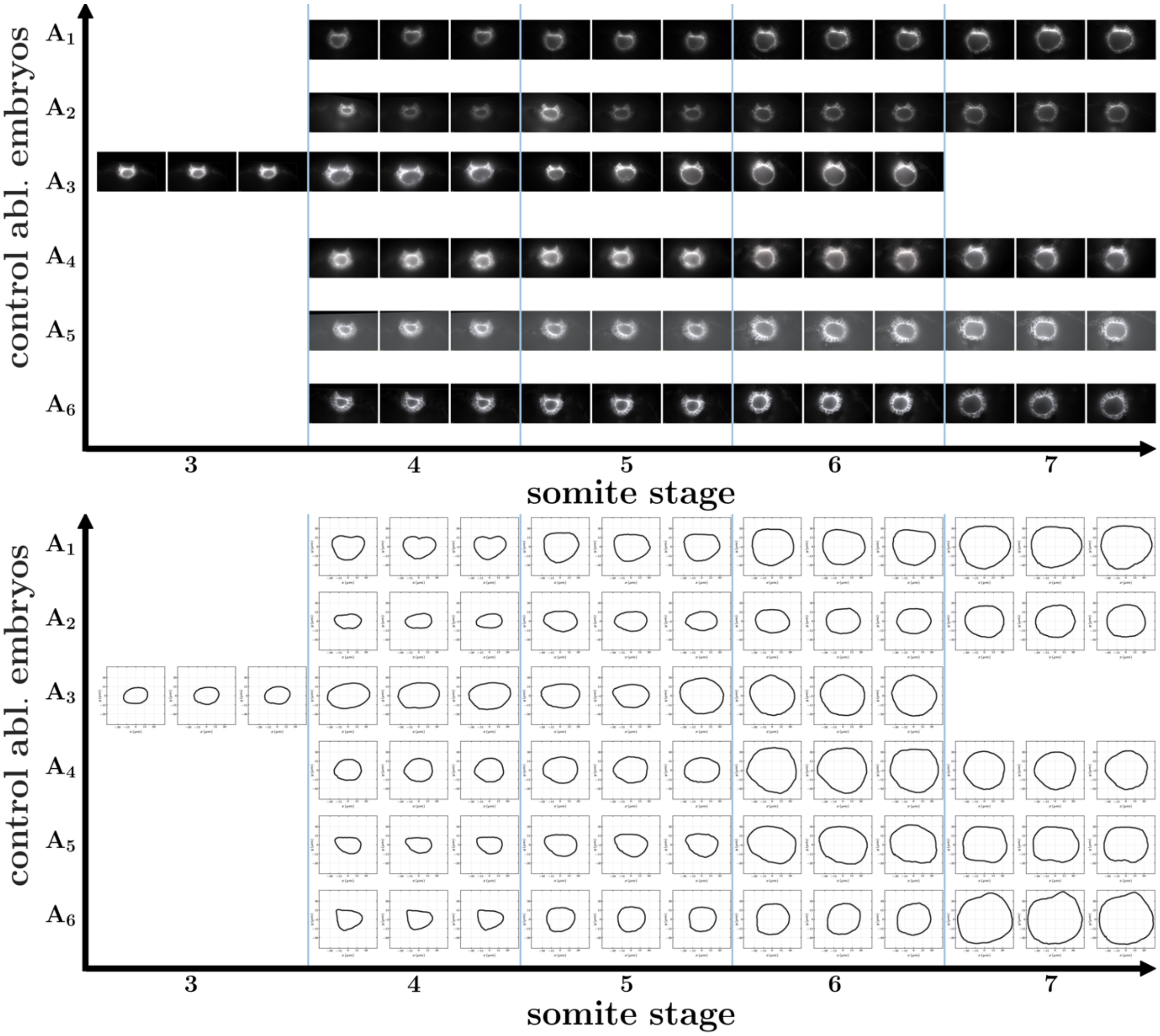


**Fig. S9. KV middle plane (A) and the corresponding lumen boundary (B) for the computation of lumen shape in control ablation experiments.** Different embryos are marked by $A_1$ to $A_6$. To calculate the lumen shape at each somite stage, three time snapshots separated by 2 minutes (0.033 somite stage) are used.

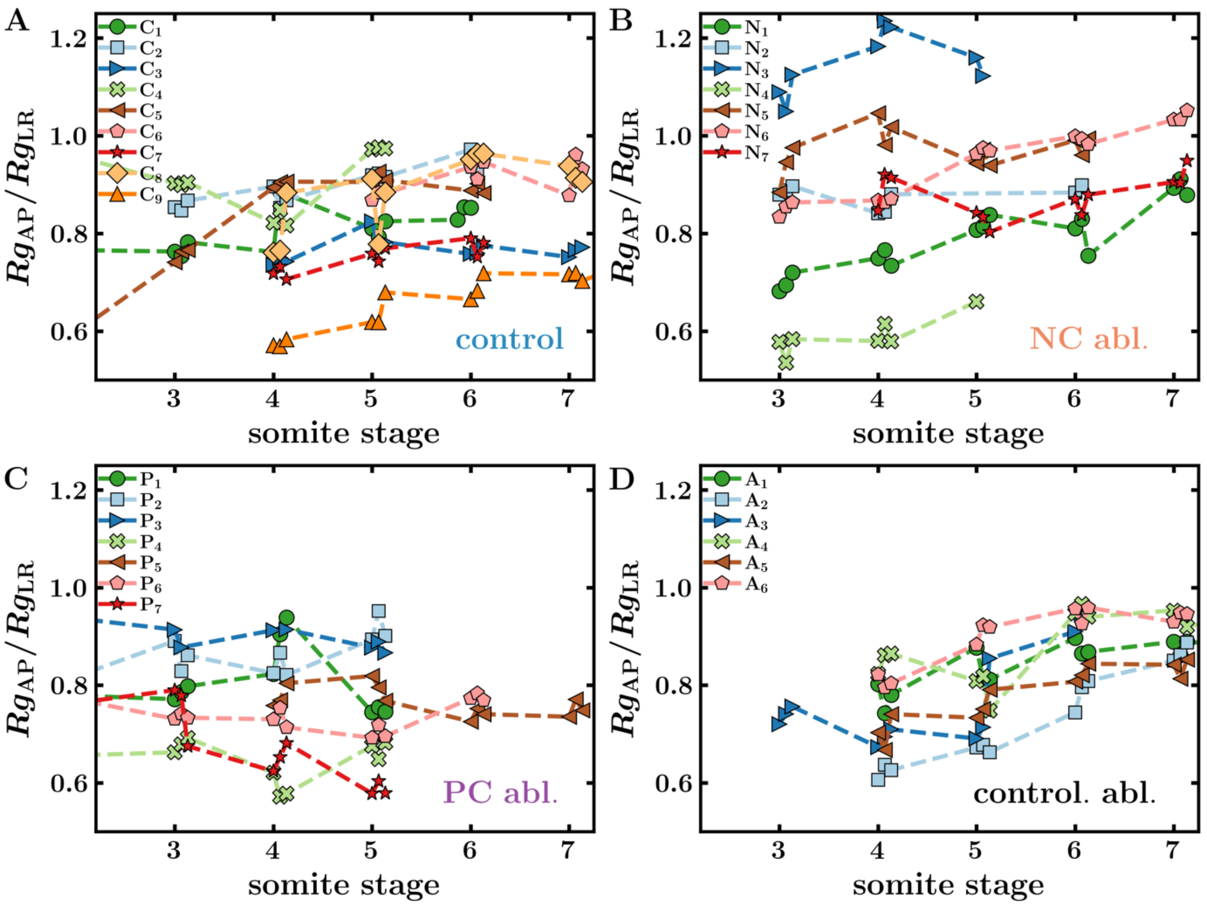


**Fig. S10. The shape of lumen in control and different ablation experiments.** The ratio of radius of gyration of lumen along AP-axis and LR-axis for individual KV as a function of somite stage in unablated control (A), notochord ablation (B), posterior cells ablation (C), and control ablation (D) experiments.

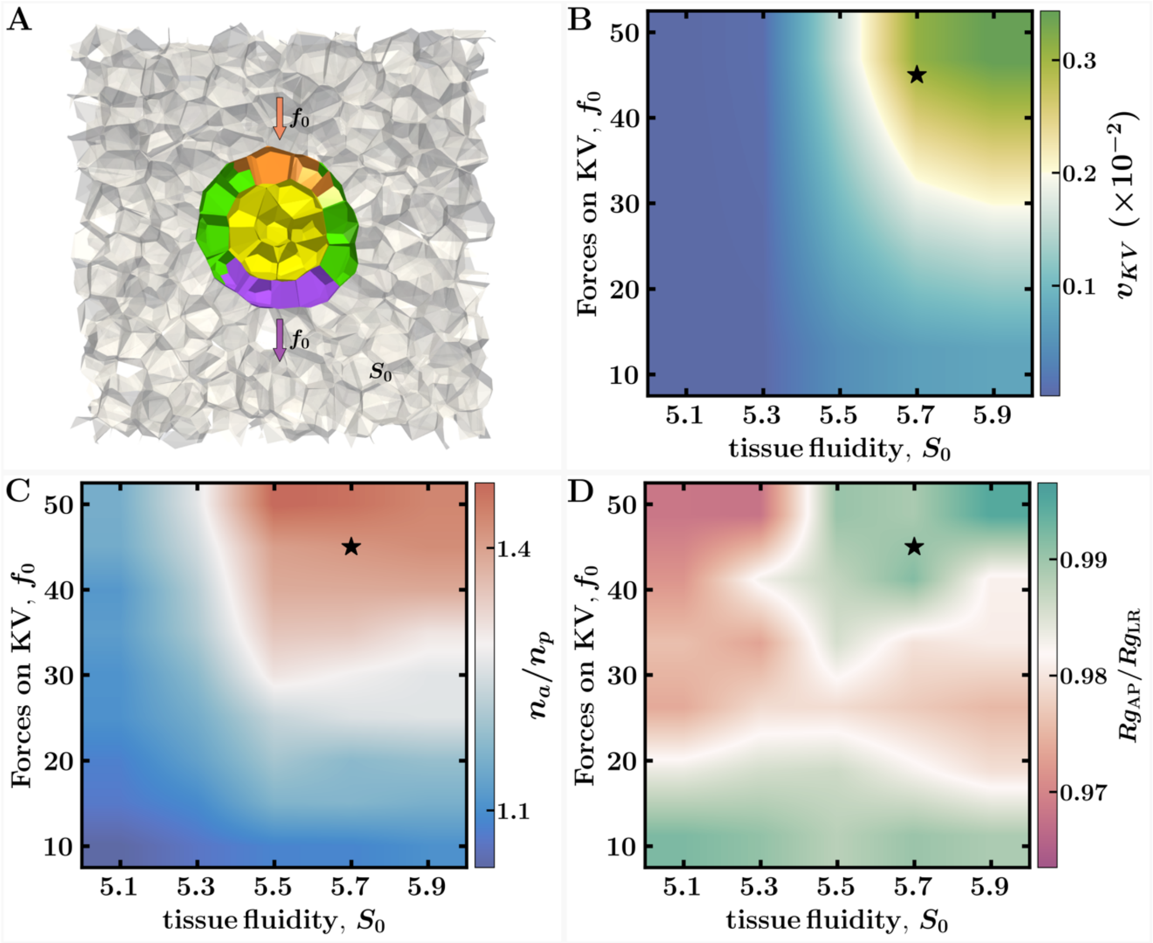


**Fig. S11. Simulations predict how KV cell shape changes depend on the tailbud tissue fluidity and forces on KV.** (A) To mimic the unablated control embryos, the pushing and pulling forces on the KV are applied to the anterior (marked by orange cells) and posterior (marked by purple cells) part of the KV. Here the same magnitude of forces, ($f_0$) are applied to the anterior and posterior region of the KV. The arrows indicate the direction of force on KV. We vary $f_0$ and the parameter that controls the tailbud fluidity in vertex models, the cell shape parameter $S_0$. Larger values of $S_0$ correspond to more fluid-like tissues (*9*). Colormap shows the KV steady state velocity (B), ratio of anterior to posterior cells (C), and the shape parameter for lumen (D) for different input values of forces on KV $f_0$ and tailbud tissue fluidity $S_0$. The star in the phase diagrams marks the parameter values that are used to generate the control case results for speed and shape changes shown in Fig. 2F, I, and J.

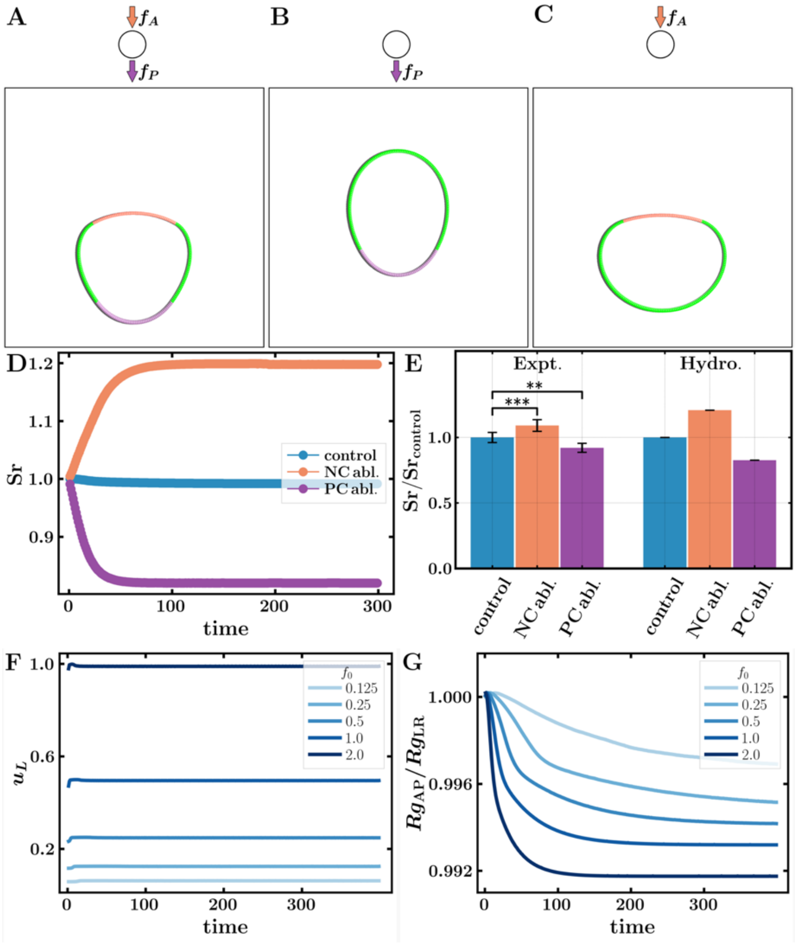


**Fig. S12 Hydrodynamic model for lumen shape changes in control and with different perturbations.** The lumen is modeled as a deformable membrane moving in a highly viscous medium. (A) To mimic the control embryos, the pushing and pulling forces on the membrane are applied to the anterior (marked by orange cells) and posterior (marked by purple cells) part of the lumen. Here the same magnitude of forces, ($f_A = f_P = 2$) are applied to the anterior and posterior region of the lumen. The arrows indicate the direction of force on lumen. For better visualization, the beads in the lumen are shown 4 times larger than their actual size. (B) To investigate the role of notochord ablation, the forces at the anterior part of the lumen are taken to be zero ($f_A = 0, f_P = 2$) (C) The forces at the posterior part of the lumen are assumed to be zero in the posterior cells ablation ($f_A = 4, f_P = 0$). (D) Temporal dynamics of the lumen shape parameter (Sr) for the three different cases. (E) A comparison of lumen shape parameter (Sr) for the three different cases between the hydrodynamic theory and experiment. (F-G) Effect of different dynamic forces on lumen speed and shape in the hydrodynamic model for control case. Here the same magnitude of forces, ($f_A = f_P = f_0$) are applied to the anterior and posterior region of the lumen. The speed of lumen (F) and lumen shape parameter (G) as a function of time for different magnitude of dynamic forces, $f_0$.

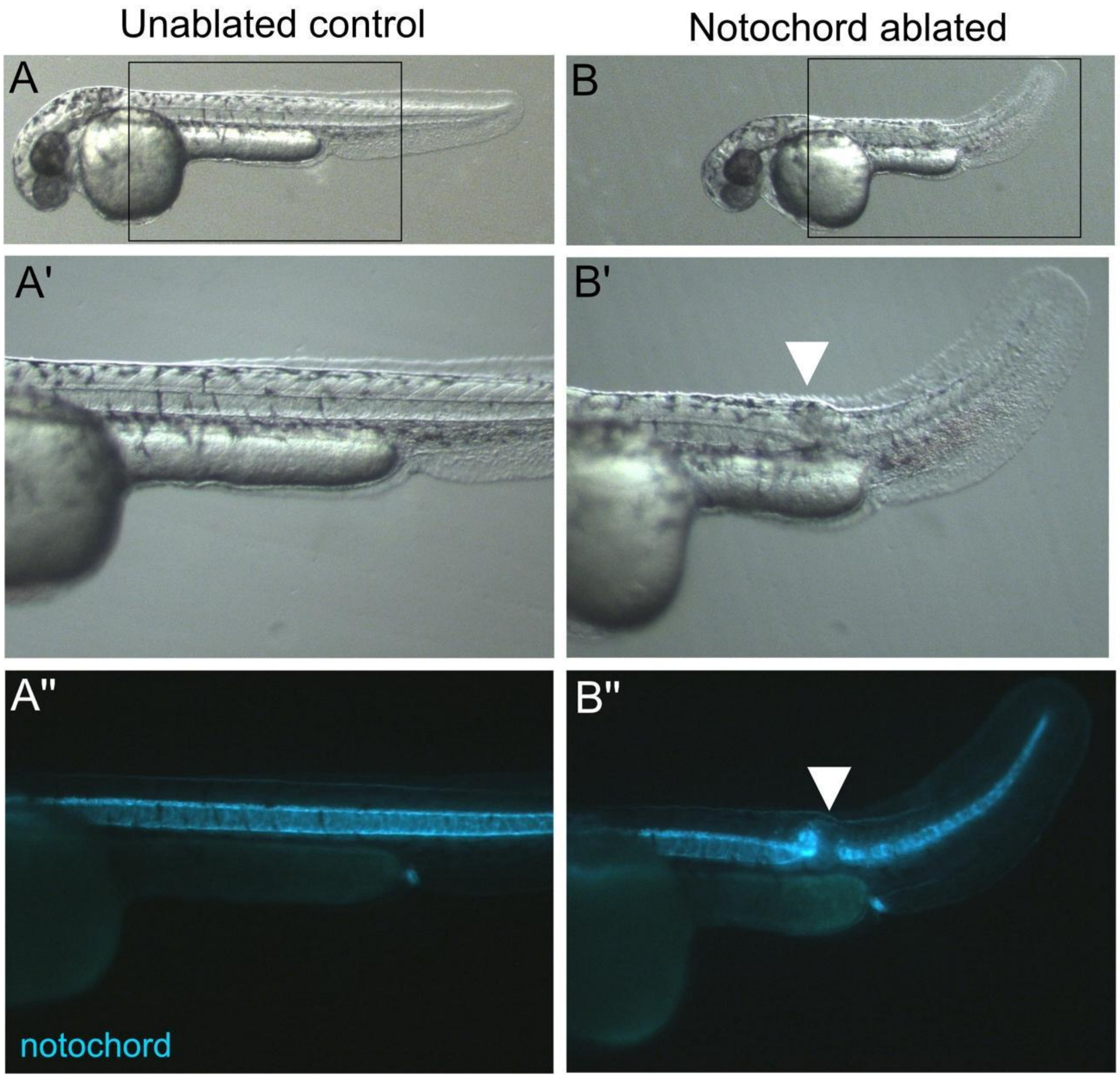


**Fig. S13. Notochord lesion persists 24 hours post ablation in notochord ablated embryos.** (A-B) Images of unablated control embryo (A) and notochord ablated embryo (B) 24 hours post ablation. (A’-B’) Higher magnification images of boxed regions in A-B. The notochord ablated embryo has a trunk lesion (arrowhead in B’) and curved tail. (A’’-B’’) Fluorescent image of embryos in A’-B’. Expression of the *Tg(-2.4shha-ABC:GFP)* transgene marks notochord cells with GFP (green). The notochord is continuous in the unablated control embryo (A’’), whereas the notochord remains severed in the notochord ablated embryo (arrowhead in B”).

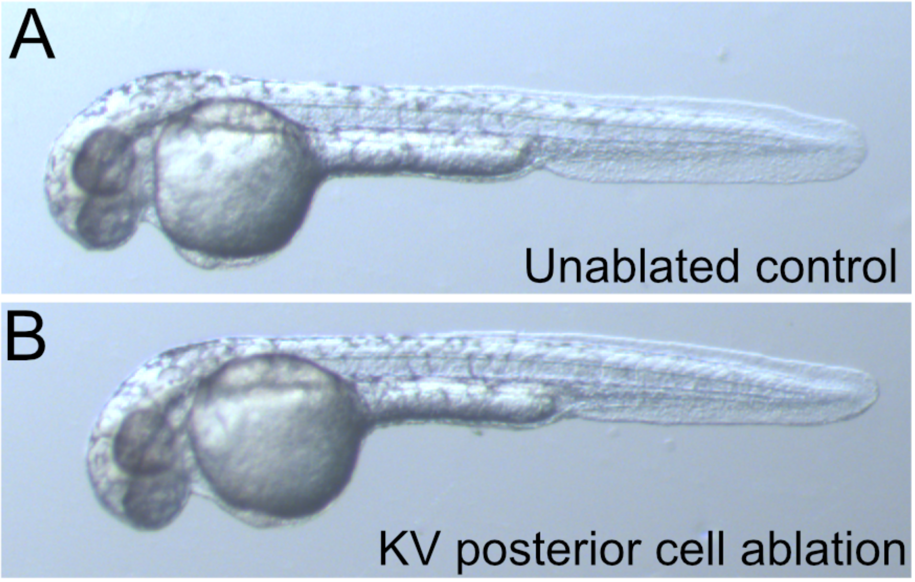


**Fig. S14. Ablation of KV posterior cells does not result in gross morphologic defects**.
(A-B) Images of unablated control embryo (A) and KV posterior cell ablated embryo (B) 24 hours post ablation. The ablation of posterior KV cells does not alter embryo gross morphology.

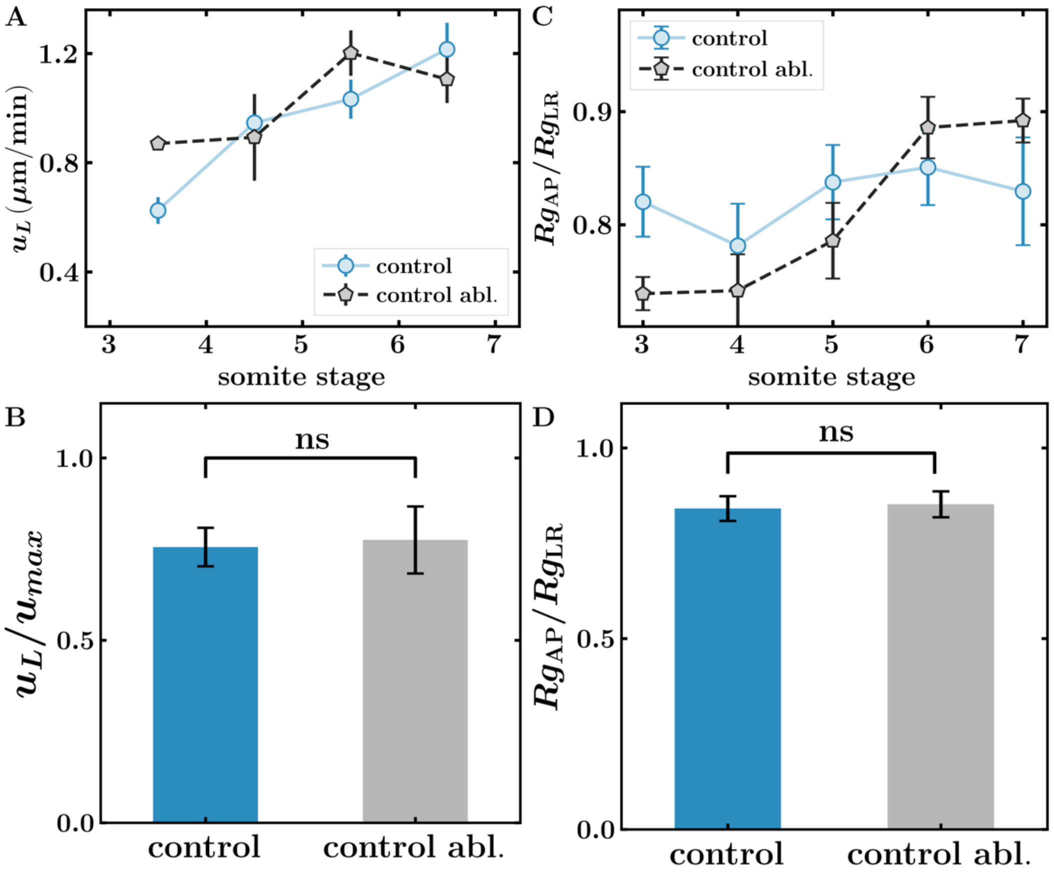


**Fig. S15. Comparison of KV speed and shape between unablated control and control ablation experiments.** (A) The average speed of KV as a function of somite stage in unablated control (n=9) and control ablation (n=6) embryos. (B) KV speed in unablated control and control ablation embryos averaged over 5-7 ss. (C) Quantification of KV shape parameter as a function of somite stage in unablated control and control ablation embryos. (D) The shape parameter averaged over 5-7 ss. ns=no significant difference.

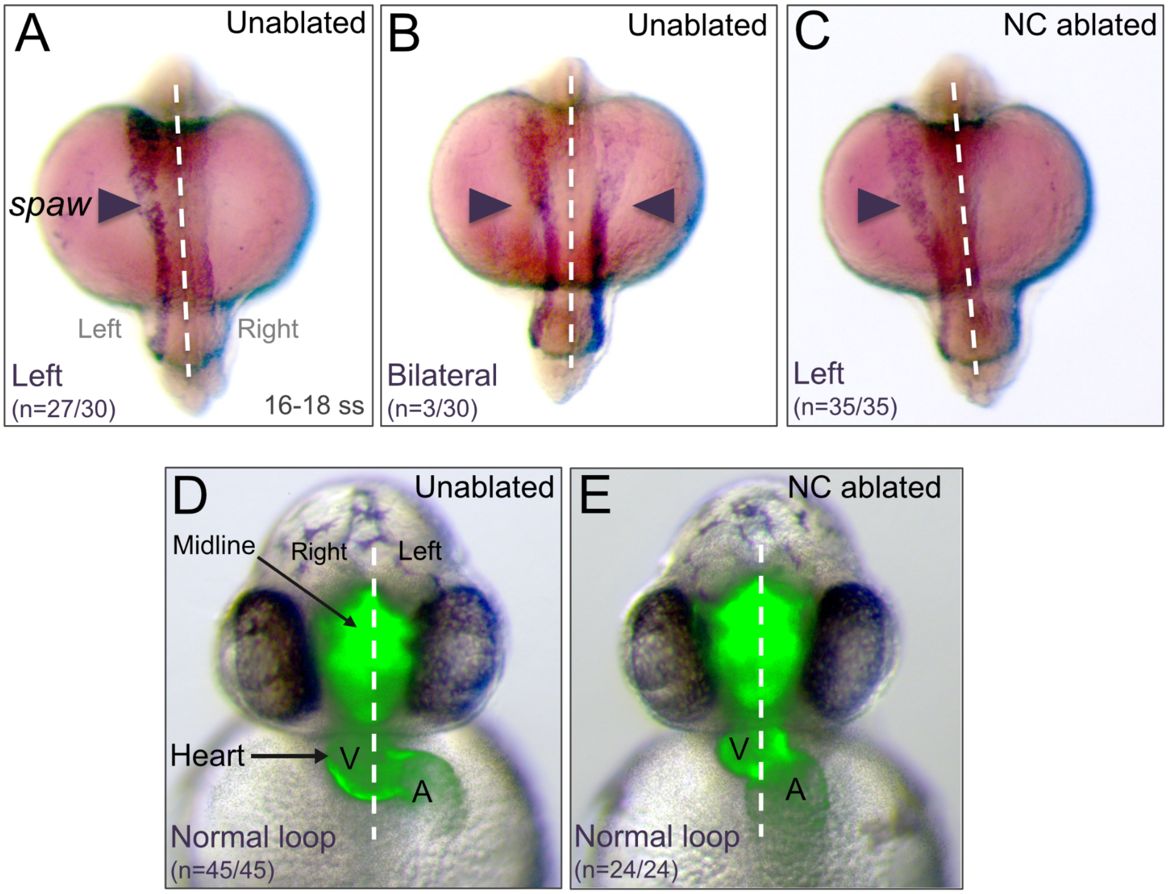


**Fig. S16. Left-right patterning is normal in notochord ablated embryos.** (A-C) RNA in situ hybridization analysis of the LR patterning marker southpaw (spaw) (a zebrafish Nodal homolog) that is asymmetrically expressed in lateral plate mesoderm at the 16 to 18 somite stages. The majority of unablated controls (n=27/30 embryos) had normal left-sided spaw expression (A) and in some cases controls had bilateral spaw (n=3/30) (B). All notochord ablated embryos analyzed (n=35/35) had normal left-sided spaw expression (C) See also Table S2. (D-E) Visualization of LR asymmetric heart looping in live embryos at 2 days post-fertilization. GFP expression labels the axial midline and the heart. Similar to unablated controls (D) all notochord ablated embryos (E) had normal rightward heart looping. White dashed lines indicate the embryo midline. V=ventricle; A=atrium of heart.

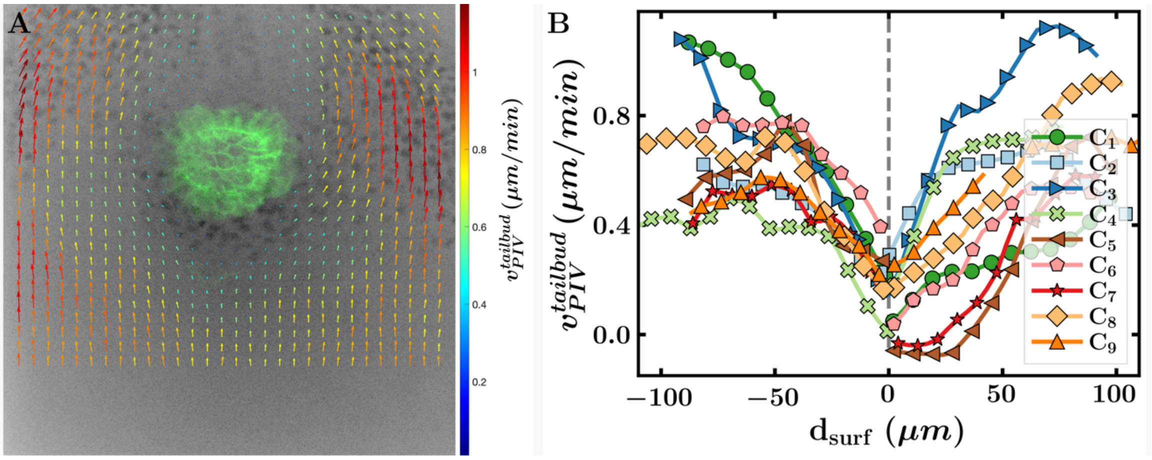


**Fig. S17. Velocity profiles of tailbud cells relative to KV motion using PIV analysis.** (A) Velocity profiles of tailbud cells relative to KV velocity in control experiments. Arrow length is proportional to the magnitude of extracted tailbud cell velocity, with the color bar also indicating velocity magnitude. The profiles shown in (A) are obtained from the $C_3$ control embryo. (B) Tailbud cell velocity (relative to KV velocity) at the equator of the KV as a function of distance from the KV surface for control, illustrating a strong velocity gradient deep into the surrounding tailbud tissue. Different colors represent individual embryos. Here, the velocity of tailbud cells is averaged across all somite stages (3ss–7ss).

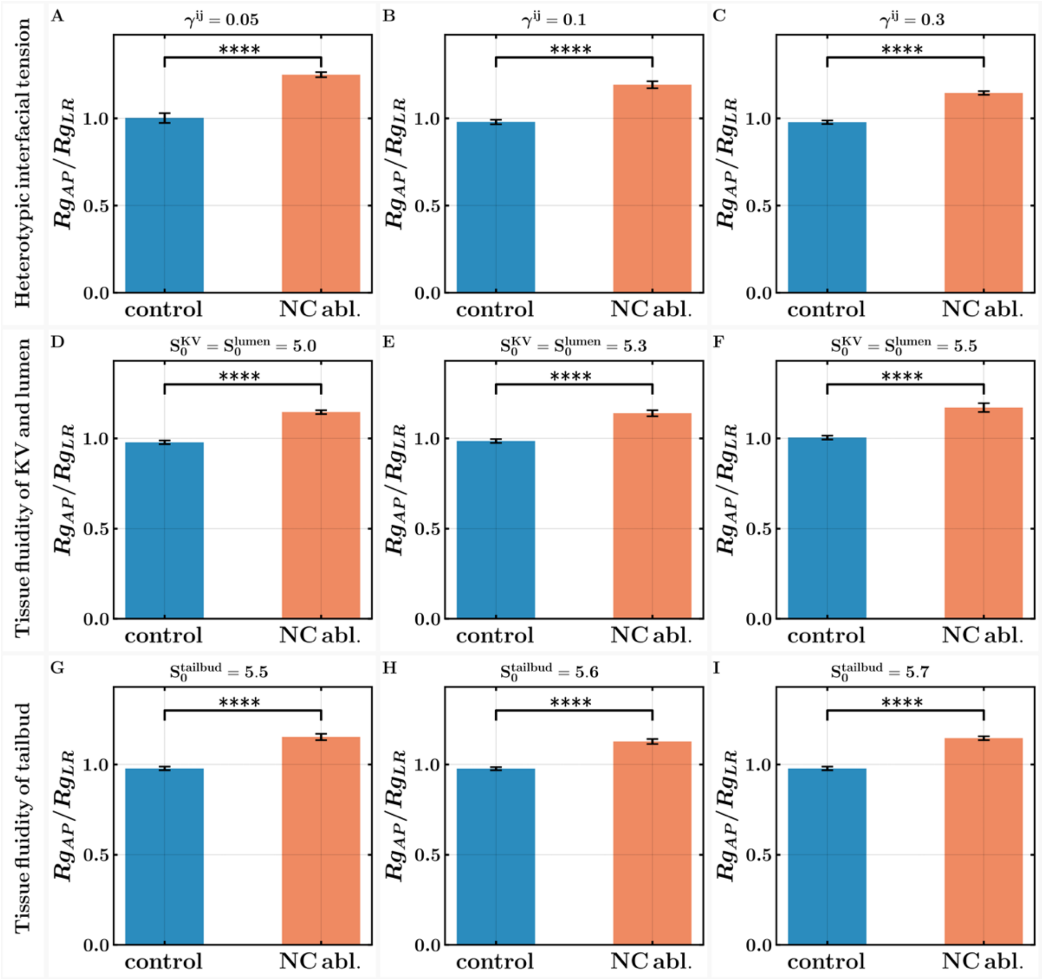


**Fig. S18. Impact of heterotypic interfacial tension and tissue fluidity on lumen shape changes in the 3D vertex model.** (A-C) The lumen shape order parameter for control and NC ablation at different heterotypical interfacial tension $\gamma^{ij} = 0.05$ (A), $\gamma^{ij} = 0.1$ (B) and $\gamma^{ij} = 0.3$ (C). Here, we keep the interfacial tension between KV and tailbud cells, $\gamma^{KV-tailbud}$ same as interfacial tension between KV and lumen, $\gamma^{KV-lumen}$. (D-F) The lumen shape order parameter for control and NC ablation at different tissue fluidity of KV and lumen $S_0^{KV} = S_0^{lumen} = 5.0$ (D), $S_0^{KV} = S_0^{lumen} = 5.3$ (E), and $S_0^{KV} = S_0^{lumen} = 5.5$ (F). Here, we take the tissue fluidity of the lumen and KV as having the same value. (G-I) The lumen shape order parameter for control and NC ablation at different tissue fluidity of tailbud $S_0^{tailbud} = 5.5$ (G), $S_0^{tailbud} = 5.6$ (H), and $S_0^{tailbud} = 5.7$ (I). Here, we take the tissue fluidity of tailbud cells in the fluid-like regime (*9*) such that KV can propel through tailbud cells (see Fig. 11). The choice of heterotypic interfacial tension and tissue fluidity does not strongly affect the lumen shape order parameter and order parameter is significantly higher for NC ablation in all cases.

### Supplemental Movies

**Movie S1: Live imaging of KV morphogenesis in the zebrafish tailbud.**
3D rendering of the zebrafish embryo tailbud in a developing *Tg(Sox17:EGFP-CAAX);Tg(ubi:mScarlet-NLS)* embryo. KV cells are labeled with membrane-localized EGFP expression (green) and all cells, including tailbud cells that surround KV, are marked by nuclear-localized mScarlet expression (white). Timestamp=Hr:Min:Sec:mSec.

**Movie S2: Movement of KV in a wild-type embryo.**
Live imaging of KV (green) and surrounding cells (grey nuclei) in a wild-type *Tg(Sox17:EGFP-CAAX);Tg(ubi:mScarlet-NLS)* zebrafish embryo between 4 ss and 7 ss. The speed of KV increases over time and lumen develops an oblate shape.

**Movie S3: Laser ablation severs the notochord.**
Live imaging of a *Tg(Sox17:EGFP-CAAX);Tg(-2.4shha-ABC:GFP)* embryo during laser ablation of a selected region of interest (green box) of notochord cells at 3 ss. Notochord cells are labeled green by GFP expression. Timestamp=Min:Sec.

**Movie S4: Live imaging suggests forces acting on KV during morphogenesis.**
Live imaging of a wild-type *Tg(Sox17:EGFP-CAAX);Tg(-2.4shha-ABC:GFP)* embryo between 3 ss and 5 ss viewed simultaneously from dorsal (left) and lateral (right) perspectives. KV cells are labeled with membrane-localized GFP expression and notochord cells are labeled by cytoplasmic GFP expression (grey). Note direct contact of the notochord with KV as it undergoes convergent extension, suggesting a pushing force on KV. In addition, posterior KV cells appear highly protrusive and could provide a pulling force. Timestamp=Hr:Min:Sec:mSec.

**Movie S5: Laser ablation of a subset of posterior KV cells.**
Live imaging of a *Tg(Sox17:EGFP-CAAX);Tg(ubi:mScarlet-NLS)* embryo during laser ablation of selected posterior KV cells (green) at 3 ss. All nuclei are labeled (grey). Timestamp=Min:Sec.

**Movie S6: Movement of KV in a notochord ablated embryo.**
Live imaging of notochord (green cytoplasm), KV (green membranes) and surrounding tailbud cells (grey nuclei) between 4 ss and 7 ss in a *Tg(Sox17:EGFP-CAAX);Tg(ubi:mScarlet-NLS);Tg(-2.4shha-ABC:GFP)* embryo after notochord ablation. The speed of KV decreases over time and the lumen expands along the AP axis to take on a prolate shape.

**Movie S7: Movement of KV in a KV posterior cell ablated embryo.**
Live imaging of KV (green) and surrounding tailbud cells (grey nuclei) between 4 ss and 7 ss in a *Tg(Sox17:EGFP-CAAX);Tg(ubi:mScarlet-NLS)* embryo after ablation of a subset of KV posterior cells. The speed of KV is not different from wild-type unablated controls, but the lumen elongates along the left-right axis and takes on a more oblate shape as compared to controls.

**Movie S8: The motion KV through tailbud cells in 3D vertex model when both forces from notochord and pulling forces from posterior cells are present (mimicking control experiment).**
The middle plane of KV is shown in the movie. The anterior and posterior part of the KV are respectively marked by orange and purple cells. The dynamic forces on the KV are $f_A = 45$, and $f_P = 45$. The tissue fluidity of tailbud cells is taken as $S_0 = 5.7$.

**Movie S9: The motion KV through tailbud cells in 3D vertex model when pushing forces from notochord are absent (mimicking NC ablation experiment).**
The middle plane of KV is shown in the movie. The anterior and posterior part of the KV are respectively marked by orange and purple cells. The dynamic forces on the KV are $f_A = 0$, and $f_P = 45$. The tissue fluidity of tailbud cells is taken as $S_0 = 5.7$.

**Movie S10: The motion KV through tailbud cells in 3D vertex model when pulling forces from posterior cells are absent (mimicking PC cells ablation experiment).**
The middle plane of KV is shown in the movie. The anterior and posterior part of the KV are respectively marked by orange and purple cells. The dynamic forces on the KV are $f_A = 90$, and $f_P = 0$. The tissue fluidity of tailbud cells is taken as $S_0 = 5.7$.

**Movie S11: The motion of the lumen, modeled as a deformable two-dimensional membrane moving in a highly viscous medium.**
(Left) The shape changes of the lumen when forces from both top and bottom are present, $f_A = 2$, $f_P = 2$ (mimicking control experiment). (Middle) The shape of the lumen when there are no forces from the anterior part of the lumen, $f_A = 0$, $f_P = 2$ (mimicking notochord ablation experiment). **(Right)** The shape of the lumen when there are no forces from the posterior part of the lumen, $f_A = 4$, $f_P = 0$ (mimicking posterior cells ablation experiment).

**Supplemental Tables**

**Table S1: Impact of notochord ablation on the anterior-posterior distribution of ciliated KV cells**

| Treatment | Embryo | Total number of ciliated cells | Number of anterior ciliated cells (Na) | Number of posterior ciliated cells (Np) | Na/Np | % of anterior ciliated cells | % of posterior ciliated cells |
|---|---|---|---|---|---|---|---|
| unablated | #1 | 54 | 34 | 20 | 1.70 | 63% | 37% |
| unablated | #2 | 45 | 25 | 20 | 1.25 | 56% | 44% |
| unablated | #3 | 66 | 40 | 26 | 1.54 | 61% | 39% |
| unablated | #4 | 66 | 36 | 30 | 1.20 | 55% | 45% |
| unablated | #5 | 58 | 38 | 20 | 1.90 | 66% | 34% |
| unablated | #6 | 61 | 34 | 27 | 1.26 | 56% | 44% |
| unablated | #7 | 28 | 16 | 12 | 1.33 | 57% | 43% |
| unablated | #8 | 39 | 24 | 15 | 1.60 | 62% | 38% |
| unablated | #9 | 39 | 26 | 13 | 2.00 | 67% | 33% |
| unablated | #10 | 62 | 35 | 27 | 1.30 | 56% | 44% |
| unablated | #11 | 42 | 27 | 15 | 1.80 | 64% | 36% |
| unablated | #12 | 49 | 29 | 20 | 1.45 | 59% | 41% |
| unablated | #13 | 61 | 37 | 24 | 1.54 | 61% | 39% |
| unablated | #14 | 68 | 45 | 23 | 1.96 | 66% | 34% |
| unablated | #15 | 67 | 36 | 31 | 1.16 | 54% | 46% |
| unablated | #16 | 40 | 24 | 16 | 1.50 | 60% | 40% |
| unablated | #17 | 47 | 31 | 16 | 1.94 | 66% | 34% |
| unablated | #18 | 40 | 25 | 15 | 1.67 | 63% | 38% |
| unablated | #19 | 55 | 31 | 24 | 1.29 | 56% | 44% |
| unablated | #20 | 49 | 33 | 16 | 2.06 | 67% | 33% |
| unablated | #21 | 38 | 21 | 17 | 1.24 | 55% | 45% |
| unablated | #22 | 24 | 14 | 10 | 1.40 | 58% | 42% |
| unablated | #23 | 35 | 20 | 15 | 1.33 | 57% | 43% |
| unablated | #24 | 57 | 38 | 19 | 2.00 | 67% | 33% |
| unablated | #25 | 62 | 41 | 21 | 1.95 | 66% | 34% |
| unablated | #26 | 51 | 33 | 18 | 1.83 | 65% | 35% |
| unablated | #27 | 67 | 38 | 29 | 1.31 | 57% | 43% |
| unablated | #28 | 24 | 14 | 10 | 1.40 | 58% | 42% |
| unablated | #29 | 48 | 31 | 17 | 1.82 | 65% | 35% |
| unablated | #30 | 40 | 27 | 13 | 2.08 | 68% | 33% |
| | Avg | 49.40 | | Avg | 1.59 | 61% | 39% |
| | SD | 13.05 | | SD | 0.30 | 4% | 4% |

| Treatment | Embryo | Total number of ciliated cells | Number of anterior ciliated cells (Na) | Number of posterior ciliated cells (Np) | Na/Np | % of anterior ciliated cells | % of posterior ciliated cells |
|---|---|---|---|---|---|---|---|
| NC ablated | #31 | 38 | 24 | 14 | 1.71 | 63% | 37% |
| NC ablated | #32 | 47 | 30 | 17 | 1.76 | 64% | 36% |
| NC ablated | #33 | 48 | 22 | 26 | 0.85 | 46% | 54% |
| NC ablated | #34 | 41 | 24 | 17 | 1.41 | 59% | 41% |
| NC ablated | #35 | 50 | 23 | 27 | 0.85 | 46% | 54% |
| NC ablated | #36 | 28 | 15 | 13 | 1.15 | 54% | 46% |
| NC ablated | #37 | 48 | 22 | 26 | 0.85 | 46% | 54% |
| NC ablated | #38 | 44 | 23 | 21 | 1.10 | 52% | 48% |
| NC ablated | #39 | 65 | 37 | 28 | 1.32 | 57% | 43% |
| NC ablated | #40 | 67 | 44 | 23 | 1.91 | 66% | 34% |
| NC ablated | #41 | 79 | 41 | 38 | 1.08 | 52% | 48% |
| NC ablated | #42 | 44 | 26 | 18 | 1.44 | 59% | 41% |
| NC ablated | #43 | 38 | 24 | 14 | 1.71 | 63% | 37% |
| NC ablated | #44 | 80 | 45 | 35 | 1.29 | 56% | 44% |
| NC ablated | #45 | 34 | 21 | 13 | 1.62 | 62% | 38% |
| NC ablated | #46 | 55 | 31 | 24 | 1.29 | 56% | 44% |
| NC ablated | #47 | 82 | 47 | 35 | 1.34 | 57% | 43% |
| NC ablated | #48 | 84 | 48 | 36 | 1.33 | 57% | 43% |
| NC ablated | #49 | 58 | 35 | 23 | 1.52 | 60% | 40% |
| NC ablated | #50 | 64 | 40 | 24 | 1.67 | 63% | 38% |

| | | | | | | | |
|---|---|---|---|---|---|---|---|
| NC ablated | #51 | 68 | 39 | 29 | 1.34 | 57% | 43% |
| NC ablated | #52 | 54 | 27 | 27 | 1.00 | 50% | 50% |
| NC ablated | #53 | 61 | 35 | 26 | 1.35 | 57% | 43% |
| NC ablated | #54 | 99 | 57 | 42 | 1.36 | 58% | 42% |
| NC ablated | #55 | 67 | 40 | 27 | 1.48 | 60% | 40% |
| NC ablated | #56 | 48 | 24 | 24 | 1.00 | 50% | 50% |
| NC ablated | #57 | 54 | 29 | 25 | 1.16 | 54% | 46% |
| NC ablated | #58 | 53 | 31 | 22 | 1.41 | 58% | 42% |
| NC ablated | #59 | 26 | 14 | 12 | 1.17 | 54% | 46% |
| NC ablated | #60 | 42 | 24 | 18 | 1.33 | 57% | 43% |
| NC ablated | #61 | 37 | 25 | 12 | 2.08 | 68% | 32% |
| NC ablated | #62 | 36 | 18 | 18 | 1.00 | 50% | 50% |
| NC ablated | #63 | 58 | 37 | 21 | 1.76 | 64% | 36% |
| NC ablated | #64 | 39 | 21 | 18 | 1.17 | 54% | 46% |
| | Avg | 54.00 | | Avg | 1.35 | 57% | 43% |
| | SD | 17.18 | | SD | 0.31 | 6% | 6% |

Supplemental Table 1 (continued): Impact of notochord ablation on the anterior-posterior distribution of ciliated KV cells.NC=notochord; Avg=average; SD=one standard deviation.

**Table S2: Impact of notochord ablation on spaw LR asymmetry**

| | spaw | expression | at 16-18 ss | | |
|---|---|---|---|---|---|
| Treatment | Left-sided | Right-sided | Bilateral | Absent | # of embryos |
| Unablated | 90% | 0% | 10% | 0% | 30 |
| NC ablated | 100% | 0% | 0% | 0% | 35 |

% of embryos pooled from four independent experiments
NC=notochord

**Table S3: Impact of notochord ablation on heart laterality**

| | Heart looping | | | |
|---|---|---|---|---|
| Treatment | Rightward | Leftward | No loop | # of embryos |
| Unablated | 100% | 0% | 0% | 45 |
| NC ablated | 100% | 0% | 0% | 24 |

% of embryos pooled from six independent experiments
NC=notochord

**Table S4: Elastic parameters in 3D vertex simulation**

| Cells | Number | $K_V$ | $K_S$ | $V_0$ | $S_0$ |
|---|---|---|---|---|---|
| Tailbud | 2048 | 10 | 1 | 1 | 5.0-5.9 |
| KV | 50 | 10 | 1 | 1 | 5.0 |
| Lumen | 17 | 10 | 1 | 1 | 5.0 |

**Table S5: Heterotypic interfacial tension between different cell types used in 3D vertex simulation**

| $\gamma^{ij}$ | Values |
|---|---|
| Tailbud-KV | 0.3 |
| KV-Lumen | 0.3 |
| Tailbud-Lumen | 2 |

**Table S6: The magnitude of dynamical forces in 3D vertex simulation**

| Figure | $f_A$ | $f_P$ |
|---|---|---|
| Fig. 2F (top), 2I (control) and 2J (control) | 45 | 45 |
| Fig. 2F (middle), 2I (NC abl.) and 2J (NC abl.) | 0 | 45 |
| Fig. 2F (bottom), 2I (PC abl.) and 2J (PC abl.) | 90 | 0 |
| Fig. 3E (control) | 45 | 45 |
| Fig. 3E (NC abl.) | 0 | 45 |

**Table S7: Simulation parameters for hydrodynamic model for lumen**

| Parameters | Values |
|---|---|
| Number of beads ($N$) | 200 |
| Viscosity ($\eta$) | 1/6 |
| Strength of WCA potential ($\epsilon$) | 0.1 |
| Area-stretching modulus ( $K_A$) | 1.0 |
| Spring constant (k) | 100 |
| Bending stiffness ($\kappa$) | 2.6 |
| Diameter of bead (2b) | 1 |
